# TriMouNet: An Algorithm for Inferring Level-1 Phylogenetic Networks from Multi-Locus Gene Tree Distributions

**DOI:** 10.64898/2026.02.14.705539

**Authors:** Qiyun Mao, Stefan Grünewald

## Abstract

With the availability of full genomes of an ever-increasing number of species, many recent phylogenetic analyses have focused on datasets with thousands of loci. In the presence of incomplete lineage sorting (ILS), many gene trees will be discordant with the species tree, which can then be estimated with a supertree method. In a separate stream of research, the reconstruction of phylogenetic networks, which aim to detect and visualize reticulations in addition to the dominant phylogenetic tree, has become common practice. Level-1 networks have the feature that every node can be interpreted as an ancestor of a subset of the taxa of interest and no two distinct cycles share a common node. TriLoNet is a method that constructs a level-1 network on all taxa from 3-taxa networks, so-called trinets. This approach is similar to modern supertree methods, but the trinets are assigned based on a single sequence alignment. Here we present TriMouNet (<u>Tri</u>net <u>M</u>ultil<u>o</u>c<u>u</u>s <u>Net</u>work), which uses the tree topology and branch length distribution in gene trees of a multilocus dataset to infer best-fitting trinets, together with scores quantifying their statistical support. These trinets are then puzzled together into a network on all taxa in a TriLoNet fashion. Experiments on simulated and real datasets show that TriMouNet can identify reticulations with low false positive rate, if the gene trees are accurate. On the other hand, TriLoNet applied to the concatenation of all loci, tends to predict wrong reticulations as the consequence of violations of model assumptions.

## Introduction

Within the last few decades, it has become common practice that phylogenetic analyses do not only attempt to construct the best-fitting phylogenetic tree but also consider reticulate evolution. This is often done by adding a phylogenetic network on the taxa of interests. Various kinds of networks and reconstruction methods have been suggested. In the article that presented the software package Splitstree4 (Huson & Bryant, 2006), Huson and Bryant suggested that “any network in which taxa are represented by nodes and their evolutionary relationships are represented by edges” might be called a phylogenetic network, and they classified several types of networks. Especially, distinguishing explicit and implicit networks has become standard terminology, and the method development for both is rather different. Explicit networks have the advantage that all nodes can be interpreted as observed taxa or as ancestors of some of the taxa, and all edges represent lineages that existed at some time or reticulation events. On the other hand, implicit networks use nodes and edges to visualize splits or clusters of the taxa (Hejase et al. 2016).

Generally, biologists prefer explicit networks because of their easier interpretability, but there is the danger to infer information that is not supported by the data. Especially the violation of common assumptions for sequence evolution, for example site independence or base composition in equilibrium, often do not introduce a strong bias favoring some trees. However, for network reconstruction these violations are likely to cause wrongly predicted reticulations (Philippe et al. 2011; Huang et al. 2022).

One of the classes of explicit phylogenetic networks that existing programs have used are Level-1 networks (Wang et al. 2001). They allow only one reticulation per biconnected component of a network within a network and can be seen as a compromise between a rooted tree and a general network that might be hard to read and not feasible to optimise for realistic data sizes. TriLoNet (Oldman et al, 2016) is a triplet-based method that first computes small networks on every set of three taxa and then merges all such ‘trinets’ (Huber and Moulton 2012) into a big network on all taxa.

While the merging process is done in a convincing way, the trinets are constructed based on single sequence alignments, and several unrealistic assumptions need to be made in order to correctly assign a trinet. Indeed a 3-way alignment cannot contain parsimony-informative site patterns, and the number of singleton sites will mainly depend on the length of the edges of a 3-taxa tree. Mild deviations from a molecular clock assumption will make it impossible to identify the strongest triplet or even a correct network.

Here we present TriMouNet (Trinet Multilocus Network), a method that assigns trinets to all sets of three in-groups and merges them into a level-1 networks in a TriLoNet-like fashion. Since the trinets come with significance scores, the merging process avoids tie-breaking.

Multilocus datasets make it possible to reconstruct many gene trees, and supertree methods like Astral (Mirarab et al. 2014) will merge them into a species tree, assuming that only incomplete lineage sorting (ILS) is responsible for gene tree discordance. ILS is modelled by the multi-species coalescent (MSC) (Degnan et al. 2009). In the presence of reticulations, the distribution of the edge lengths and topologies of the gene trees will deviate from the MSC, and if an outgroup is included this distribution makes it possible to infer trinets.

Huber and Moulton characterized all trinets that can be induced by level-1 network and found 8 binary types. Here we model reticulations as instantaneous events, following ideas by Moret et al. (2004). As a consequence, reticulation edges are drawn horizontally and we distinguish only 6 classes of trinets as shown in Figure 1. For S_1_ trinets, the distribution of the edge lengths and topologies of the gene trees allow us to identify which taxon is the descendant of the reticulation node; in S_2_ and N-type trinets, this direction cannot be determined from the data. Therefore, a reticulation edge is directed for S_1_ and undirected else. Notably, the four NT-type trinets in Figure 1 can be obtained from T_1_ by replacing one or both bifurcations by a reticulate binet S_0_.

**FIG. 1.**
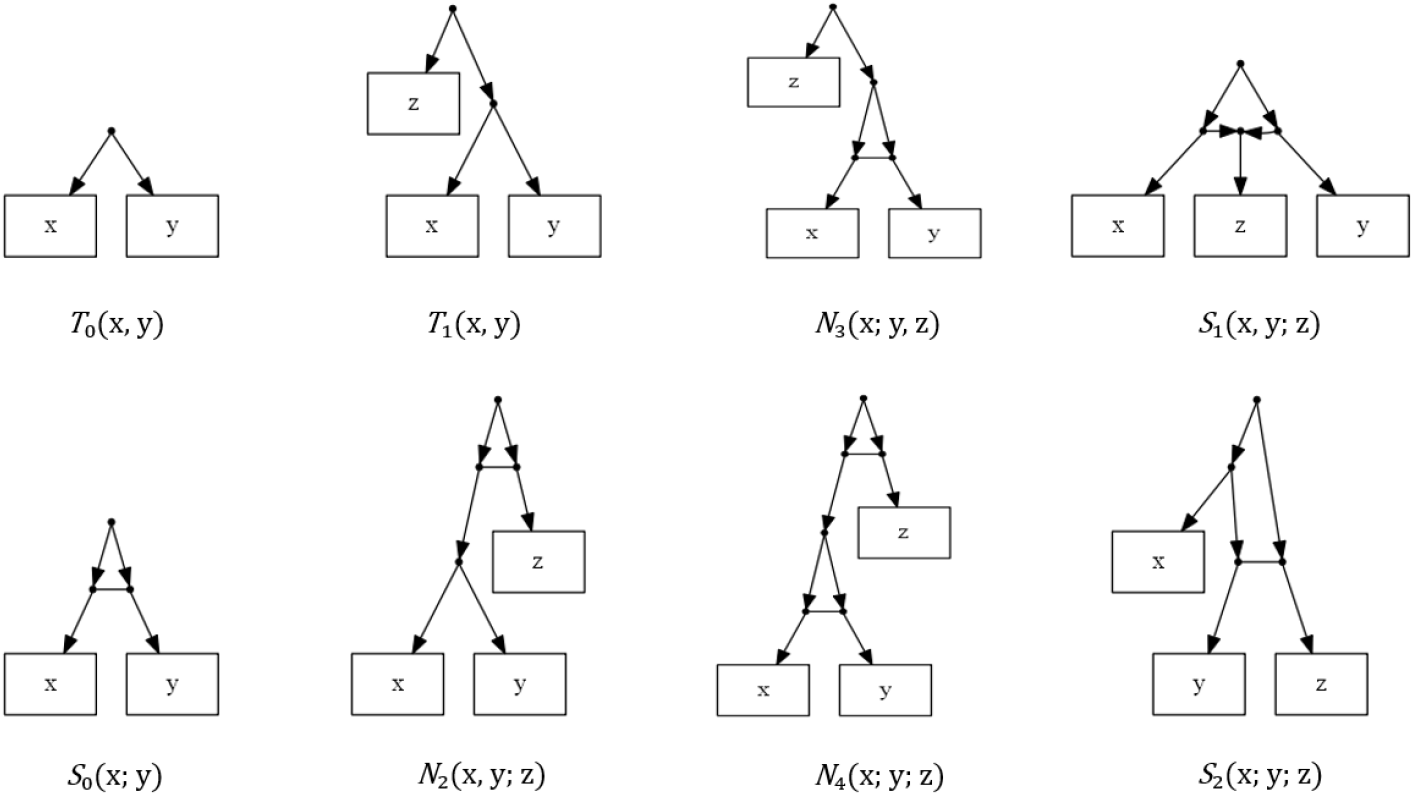
The two types of binets and the six types of trinets.

In the absence of reticulations, the MSC assumes that the probability for two lineages to not coalesce within time t (in coalescence units) equals *e*^−*t*^. Given a 3 taxa species tree with sister taxa a and b and outgroup c and with interior edge length t, the probability that the triplet ab|c is observed in a gene tree is 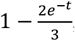, while the probabilities for ac|b and bc|a both are 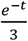. Since both triplets discordant with the species trees have the same probability, the numbers of gene trees displaying the second most frequent and the most rare triplet should be close to equal. If the observed counts for these triplets are significantly different, then a reticulation is likely. This asymmetry has been used to detect reticulations frequently for example for SNP data (Green et al. 2010) and retrotransposable elements (Springer et al. 2020). We apply a binomial test to quantify the significance.

If unique triplet is likely to be correct, we assign one of the types T_1_, N_2_, N_3_, N_4_. Given a rooted triplet ab|c with outgroup o, we quantify edge-length asymmetry using two scale-free ratios: the cherry ratio V_1_ (normalized ab distance within triplet ab|c) and the pending ratio V_2_ (mean normalized distance from c to each cherry taxon within triplet ab|c). We then use the empirical distributions of (V_1_, V_2_) across loci to select the final trinet topology.

If two different triplets are likely to be partially correct, we assign one of the types S_1_ and S_2_. We then use an additional scale-free summary statistic, V_3_, defined as the ratio of the common-edge length to the total length along the path from the common taxon to the outgroup, and use the empirical distribution of V_3_ across loci to distinguish S_1_ from S_2_.

## Results

### Simulated Data

#### TriMouNet analysis of simulated trinets

To evaluate the performance of the trinet assignment of TriMouNet, we conducted simulations for all trinet types.The results for types N_2_, N_3_ and N_4_ are shown in Figure 2. For these trinets, a mixture distribution is correct for different types of edge length ratio: the pending ratio in N_2_, the cherry ratio in N_3_, and both ratios in N_4_.

**FIG. 2.**
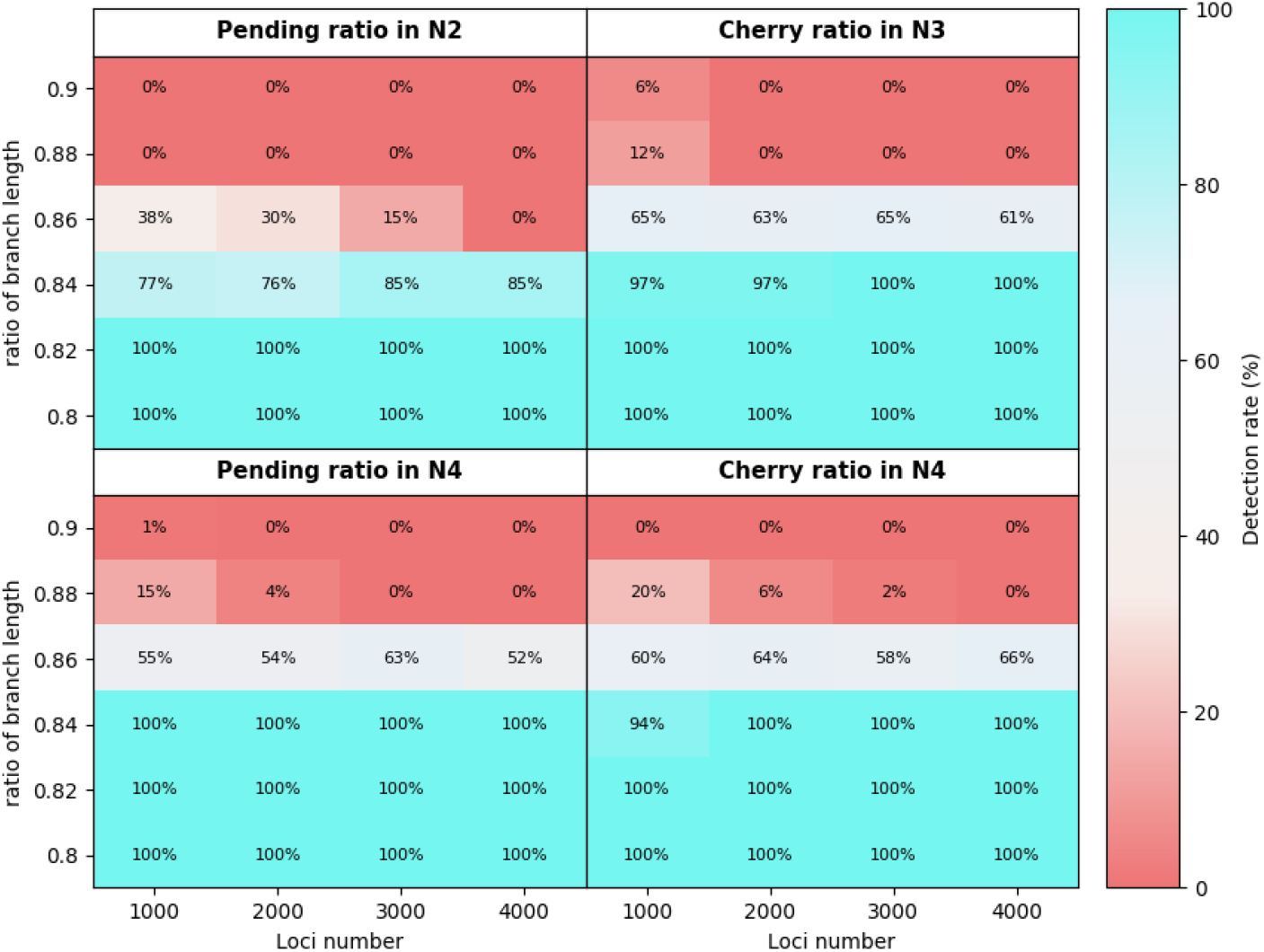
Heatmap of accuracy of mixture detection across trinet topologys.

For each simulation, we used msprime (Baumdicker et al. 2022), a coalescent-based simulator implemented in Python, to generate 100 independent multilocus datasets under controlled conditions. The number of loci varied from 1000 to 4000, and the ratio of branch lengths associated with the cherry edge or the pending edge was systematically varied from 0.8 to 0.9, with smaller ratios corresponding to stronger reticulation signals. We first reconstructed gene trees using IQ-TREE 2 (Bui Quang Minh et al. 2020) with default settings, then we computed the statistics |Lv_1,1_-Lv_1,2_| for cherry edges and |Lv_2,1_-Lv_2,2_| for pending edges (see Methods section for definitions). A typical distribution of an edge length ratio (trinet N_2_, ratio of branch length: 0.8, 4000 loci) is shown in Figure 3. For this easy example, the distribution is clearly bimodal, while for higher ratios of branch length, the two peaks cannot always be easily detected.

**FIG. 3.**
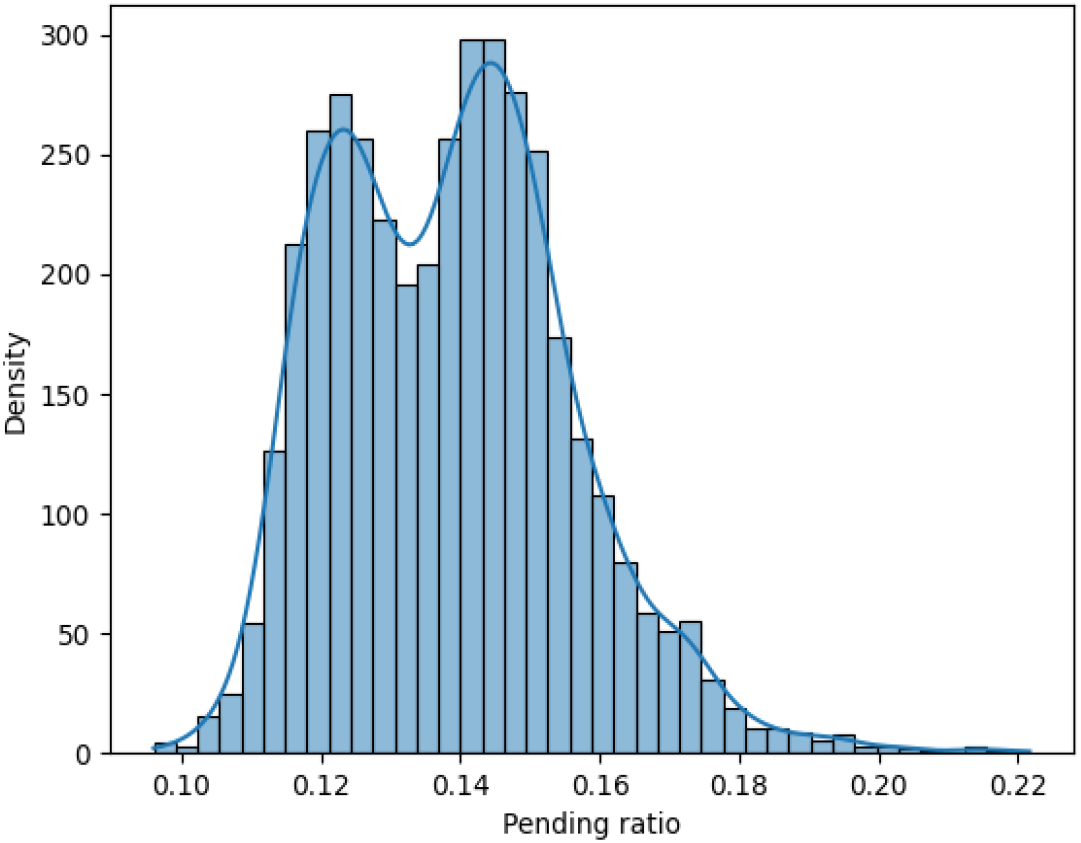
The distribution of the pending ratio for N_2_ for a simulated dataset with 4000 loci and a ratio of branch length of 0.8.

The heatmaps summarize the detection rates across the 100 replicates. Specifically, each cell represents the proportion of datasets in which the corresponding statistic exceeds a predefined threshold κ, where κ∈ {8,16,24,32} depending on the column. The rationale for setting κ proportional to the number of loci (specifically, κ= (number of loci)/125) is provided in the Supplementary Material. Warmer colors indicate lower detection rates, whereas cooler colors indicate higher detection rates.

Overall, the results demonstrate that the detectability of bimodal signals depends jointly on branch-length ratio and number of loci. As expected, detection power increases with stronger signal (smaller branch-length ratios) and larger numbers of loci, while differences among N_2_, N_3_, and N_4_ reflect the distinct placement of bimodal signals within each trinet topology.

The difference between 100% and the accuracy values in Fig. 2 corresponds to false negatives (the distribution is mixture but TriMouNet fails to detect it). False positives do not occur for the cherry ratio in N_2_ or the pending ratio in N_3_ because the threshold is chosen to be conservative enough to avoid predicting wrong reticulations for these data sizes (see Supplementary Material for details).

#### TriMouNet analysis of simulated networks

To evaluate the performance of TriMouNet under controlled simulation conditions, we followed a three-step simulation–reconstruction–inference pipeline. First, we specified an underlying reticulate network containing three reticulation nodes, which gives rise to eight distinct guiding trees corresponding to all possible combinations of parental inheritance. Multilocus sequence data were simulated using msprime across the eight guiding trees. For each tree, we generated 500 loci, resulting in a comprehensive dataset of 4,000 loci. Second, for each simulated locus, we reconstructed a gene tree using IQ-TREE 2. Finally, the collection of reconstructed gene trees was used as input to TriMouNet to infer a phylogenetic network, which was then compared against the true underlying network composed of the eight guiding trees.

Our simulated network features a reticulated cherry (taxa B, C), an internal S-type cactus (taxa F, G, H), and a basal cactus involving all taxa.

Starting from the simulated network shown in Figure 4(i), we generated a series of simulated networks with decreasing reticulation signal strength by systematically modifying coalescent times associated with reticulated cherries and the lengths of interior branches. Only the strong-signal simulated network is shown in Fig. 4; the remaining networks are described in the text. The simulated network also includes an outgroup taxon *O* with a coalescent time of 30, which is omitted from the figure for simplicity and space considerations.

**FIG. 4.**
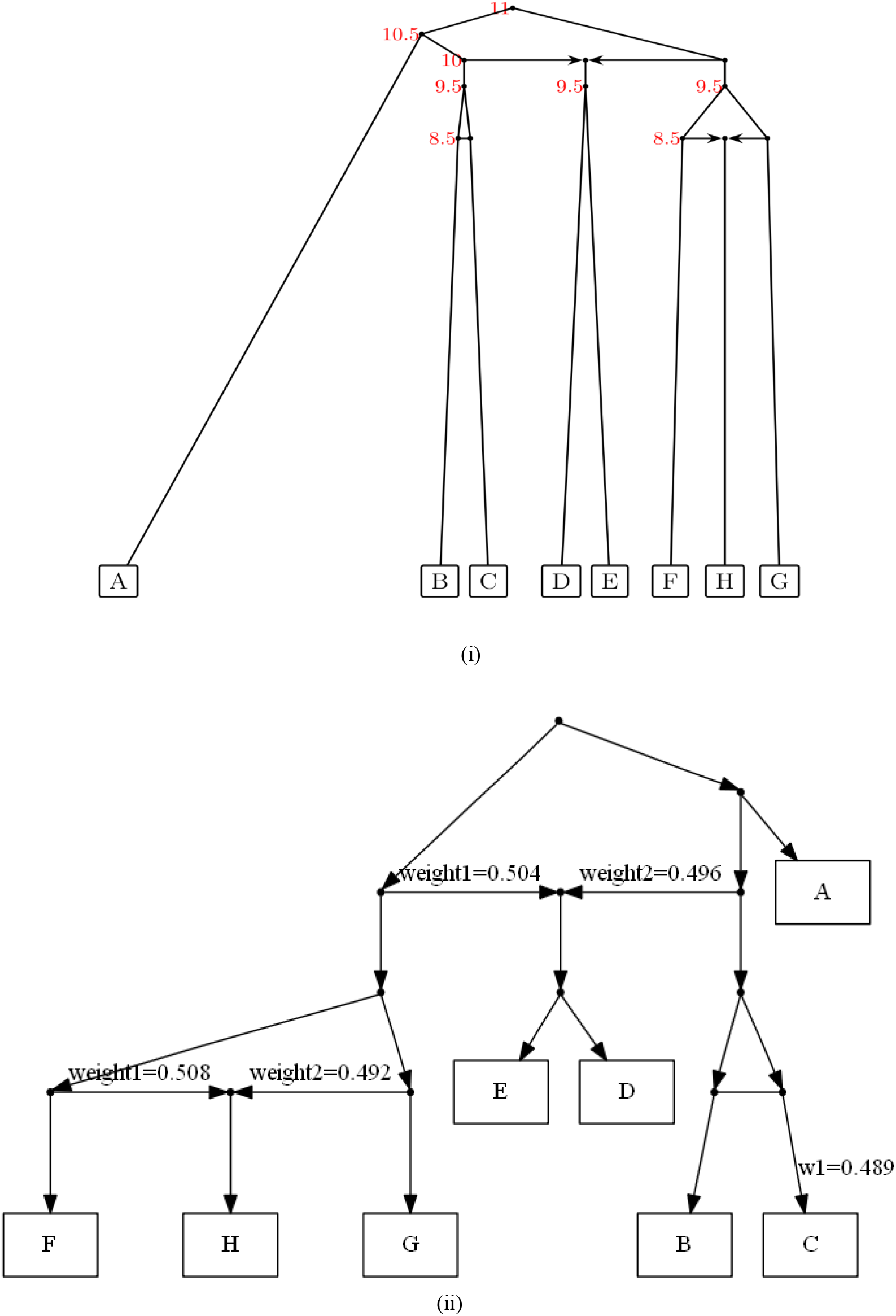
Strong-signal simulated network (i) and the corresponding network reconstructed by TriMouNet (ii).

In the strong-signal network (Fig. 4(i)), the coalescent times are well separated (9.5 vs. 8.5), and interior branches are relatively long (length = 0.5), resulting in minimal incomplete lineage sorting within our three simulated networks. Reticulation signals were progressively weakened by reducing the time difference between coalescent times and using shorter interior branches. Specifically, the intermediate-signal network uses coalescent times of 9.8 vs. 9.4 with an interior branch length of 0.2, whereas the weak-signal network uses coalescent times of 9.9 vs. 9.7 with an interior branch length of 0.1.

Figure 4(ii) shows the network reconstructed by TriMouNet under the strong-signal simulated network. In this case, TriMouNet exactly recovers the true topology, correctly identifying both the reticulated cherry and the overall cactus structure. For each reticulated cherry, our algorithm outputs a parameter w_1_, which here represents the merged support weight for the lower-level coalescent configuration within that reticulated cherry (i.e., the more recent last common ancestor indicated by the undirected edge). The exact aggregation rule used to compute w_1_ is detailed in the Methods and Supplementary Materials. All weights of the reconstructed network are close to the correct value of 50%.

Under the intermediate-signal simulated network, TriMouNet fails to detect the reticulated cherry, accompanied by an increased inferred weight of the dominant reticulation edge of the basal cactus (from 50.4% to 54.5%). Under the weakest-signal simulated network, TriMouNet neither recovers the reticulated cherry involving taxa B and C nor the correct taxon ordering within the basal cactus (taxon A and cluster (B, C) interchange), and the dominant reticulation edge further increases to 60.6%.

### Biological Data

We next applied TriMouNet to three empirical biological datasets and compared the inferred networks with both previously published phylogenetic analyses and results obtained using TriLoNet. Particular attention was given to reticulation events and splits that are inconsistent with canonical species trees.

#### TriMouNet analysis of yeast data

Yeast provides a well-established testing ground for phylogenetic workflows, supported by its deeply annotated genetic landscape and extensive evolutionary characterization (Hittinger 2013). We analyzed 1,233 single-copy amino acid loci from the Shen et al. (2016) yeast dataset, selecting *S. cerevisiae* and 10 relatives as ingroups and *Candida albicans* as outgroup. We used both TriMouNet and TriLoNet, directly comparing the inferred splits and reticulation nodes obtained from the same multilocus input.

Figure 5(i) illustrates the phylogenetic network inferred by TriLoNet from the concatenated amino acid alignment. In this topology, *Saccharomyces kudriavzevii* is positioned beneath a reticulation node, a placement that is also recovered by TriMouNet (Fig. 5(ii)). However, beyond this shared signal, TriLoNet collapses *S. kudriavzevii, S. mikatae, S. cerevisiae, S. paradoxus* and the cherry (*S. eubayanus, S. uvarum)* into a single cactus, missing the well-established cluster comprising *S. cerevisiae, S. paradoxus* and *S. mikatae*. In addition, both the Tri-MouNet-inferred network and the species tree recovered by Shen et al. (2016) but not TriLoNet resolve the *Saccharomyces* species as a sister group to

**FIG. 5.**
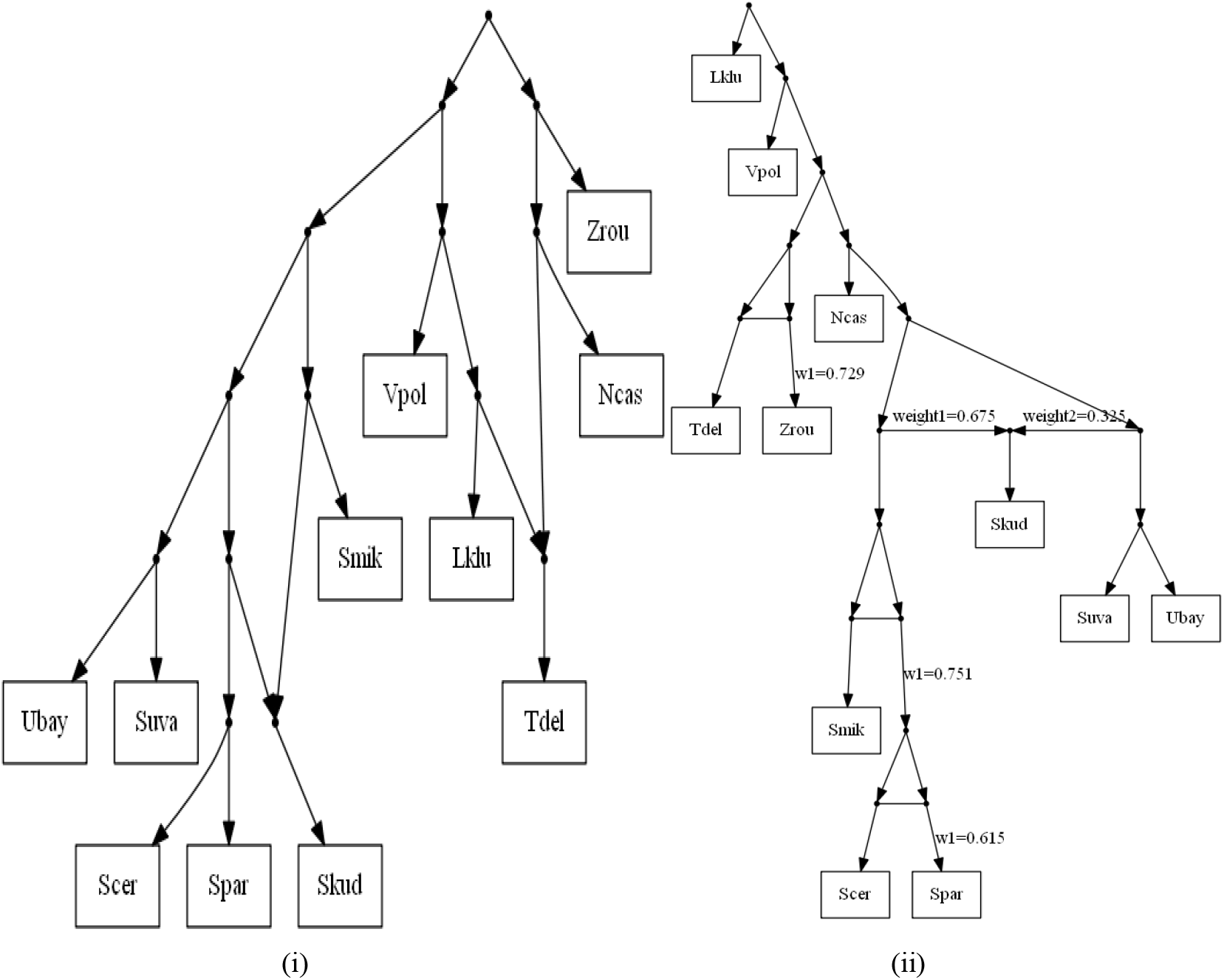
Phylogenetic network of yeast based on 1,233 single-copy amino acid loci inferred by TriLoNet and TriMouNet. Taxon abbreviations used in the networks are as follows: Scer, *Saccharomyces cerevisiae*; Spar, *S. paradoxus*; Smik, *S. mikatae*; Skud, *S. kudriavzevii*; Suva, *S. uvarum*; Ubay, *S. eubayanus*; Ncas, *Naumovozyma castellii*; Vpol, *Vanderwaltozyma polyspora*; Tdel, *Torulaspora delbrueckii*; Zrou, *Zygosaccharomyces rouxii*; Lklu, *Lachancea kluyveri*

##### Naumovozyma castellii

Additional differences between TriLoNet and TriMouNet are observed in the placement of Torulaspora delbrueckii and Zygosaccharomyces rouxii. These two taxa are strongly supported as sister species in previous phylogenetic analyses, including the species tree inferred by Shen—a relationship consistently recovered by TriMouNet.

In contrast, the TriLoNet-inferred network fails to resolve this split, instead positioning T. delbrueckii beneath a reticulation node while assigning Z. rouxii as a separate leaf within the cactus structure. In addition, TriMouNet but not TriLoNet recognized that *Lachancea kluyveri* is basal to all other ingroups, as it was observed by Shen et al. (2016) and others. On the negative side, TriMouNet misplaces *Vanderwaltozyma polyspora*. This taxon should form a cluster with the *Saccharomyces* genus and *Naumovozyma castellii*, because all of these taxa emerged from a whole-genome duplication. exhibits an ordering inconsistency among *Naumovozyma castellii, Vanderwaltozyma polyspora, Torulaspora delbrueckii*, and *Zygosaccharomyces rouxii*. With many published studies failed to reconstruct this cluster, including one of the analyses in Shen et al. (2016). For the trinet involving *N. castellii, V. polyspora*, and *Z. rouxii*, loci are almost evenly divided among the three possible resolutions: 280 loci group *N. castellii* with *V. polyspora*, 279 loci group *V. polyspora* with *Z. rouxii*, and 277 loci group *N. castellii* with *Z. rouxii*. As a result, the split corresponding to the species tree does not emerge as the dominant signal in the multilocus summary. Under such near-tie conditions, even subtle systematic biases can perturb the inferred coalescence order, leading to an incorrect final topology. Further inspection suggests that compositional heterogeneity and long-branch attraction may skew these marginal differences, thereby exacerbating the observed ordering ambiguity.

The TriMouNet-inferred network also identifies a reticulation event involving *Saccharomyces kudriavzevii*, supported by two alternative topological configurations. The primary topology (67.5% weight) associates *S. kudriavzevii* with the (*S. mikatae*, (*S. cerevisiae, S. paradoxus*)) clade, whereas a secondary configuration (32.5% weight) links it with (*S. uvarum, S. eubayanus*). This distribution indicates asymmetric yet substantial support for both parental histories, consistent with a complex reticulate origin for *S. kudriavzevii*. The same reticulation with slightly different taxa set has already been hypothesized by Holland et al. (2004). TriMouNet constructed 3 reticulated cherries (*S. cerevisiae, S. paradoxus*) with w_1_=0.615; *S. mikatae* and the reticulated cherry (*S. cerevisiae, S. paradoxus*) with w_1_=0.751 and (*T. delbrueckii, Z. rouxii*) with w_1_=0.729. Collectively, these values indicate that, across these reticulated cherries, the inferred signal places greater weight on the later coalescent configuration than on the history with the earlier coalescent.

#### TriMouNet analysis of Cupressaceae data

Single-copy nuclear genes are widely used as phylogenomic markers because they are putatively orthologous and reduce complications from paralogy, while providing large numbers of loci for coalescent-based inference (Li et al. 2017). Under the multi-species coalescent, loci are treated as independent genealogical replicates (no recombination within loci and free recombination among loci), so discordance across loci can be interpreted as biological signal rather than technical artifact (Degnan & Rosenberg 2009; Zhu et al. 2022). To explore the applicability of TriMouNet to single-copy nuclear datasets, we analyzed 1,811 single-copy nuclear genes sampled from 11 Cupressaceae species and *Papuacedrus papuana* as outgroup (Li et al. 2021), reconstructed a phylogenetic network using TriMouNet, and benchmarked the inferred topology against that obtained with TriLoNet from the same multilocus input (Fig. 6).

**FIG. 6.**
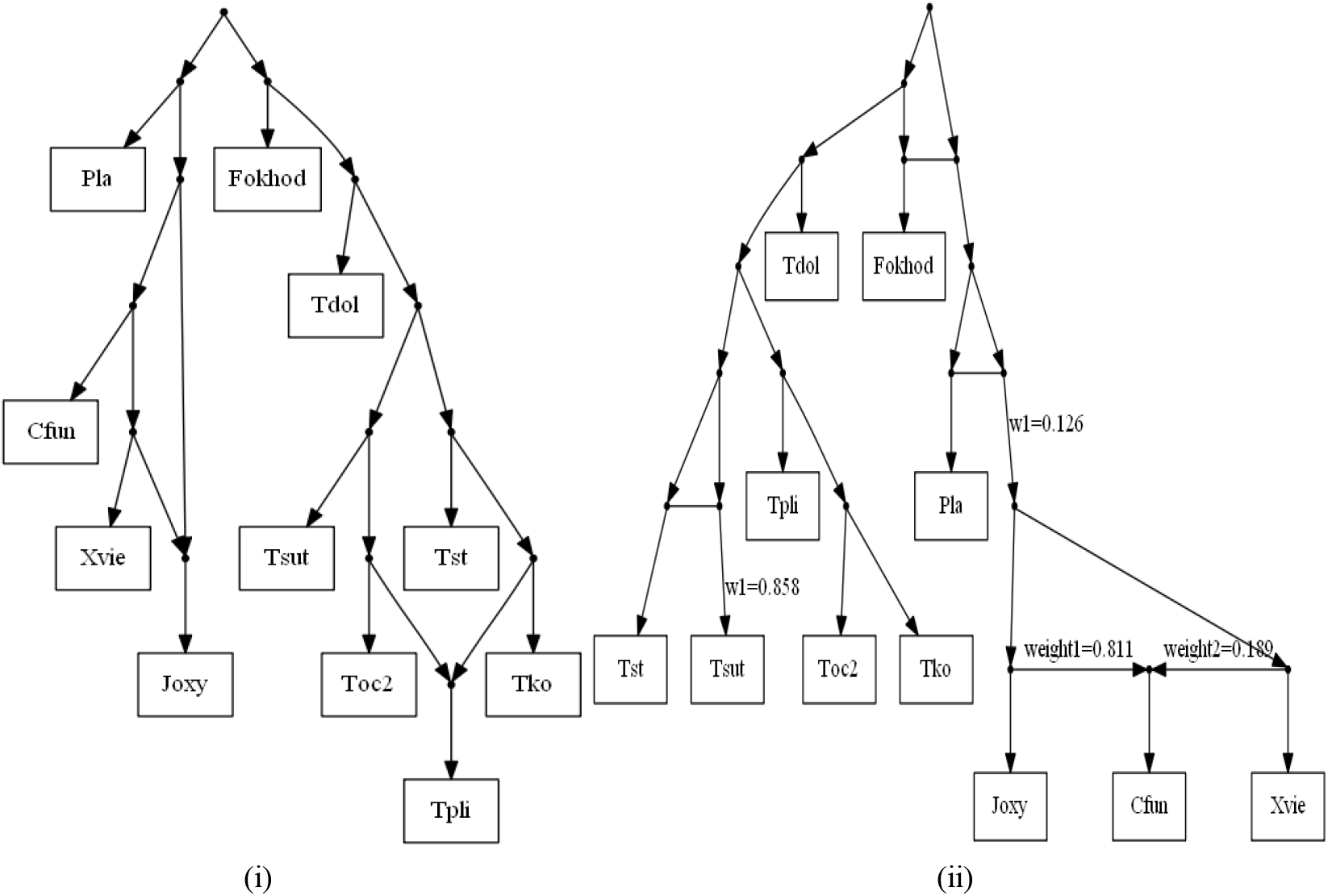
Phylogenetic network of Cupressaceae based on 1,811 single-copy nuclear genes inferred by TriLo-Net and TriMouNet. Taxon abbreviations used in the networks are as follows: Fokhod, *Fokienia hodginsii* (Fujian cypress); Pla, *Platycladus orientalis* (Oriental arborvitae); Cfun, *Cupressus funebris* (Chinese weeping cypress); Joxy, *Juniperus oxycedrus* (prickly juniper); Xvie, *Xanthocyparis vietnamensis* (Vietnamese golden cypress); Tdol, *Thujopsis dolabrata* (Hiba arborvitae); Tsut, *Thuja sutchuenensis* (Sichuan thuja); Tst, *Thuja standishii* (Japanese thuja); Tpli, *Thuja plicata* (western red cedar); Toc2, *Thuja occidentalis* (northern white cedar); Tko, *Thuja koraiensis* (Korean thuja).

Both methods recovered a well-supported clade comprising the five *Thuja* species (arborvitae)—*T. occidentalis* (northern white cedar), *T. koraiensis* (Korean thuja), *T. plicata* (western red cedar), *T. standishii* (Japanese thuja), and *T. sutchuenensis* (Sichuan thuja)—together with the closely related *Thujopsis dolabrata* (Hiba arborvitae). However, TriLoNet largely collapsed this clade into a single cactus and failed to resolve several biologically informativeinternal splits. Specifically, TriLoNet failed to recover the sister relationships between *T. occidentalis* + *T. koraiensis* and *T. standishii* + *T. sutchuenensis*, notably, T. standishii and T. sutchuenensis form a reticulated cherry with w_1_=0.858 in the TriMouNet-inferred network. Both of which have been consistently recovered in previous phylogenetic studies based on nuclear and plastid data (Li et al. 2021).Within the TriMouNet network, *Cupressus funebris* (Chinese weepingcypress), *Xanthocyparis vietnamensis* (Vietnamese golden cypress), and *Juniperus oxycedrus* (prickly juniper) constitute a small cactus. Among the inferred topologies, 81.1% support the grouping ((*J. oxycedrus, C. funebris*), *X. vietnamensis*), whereas 18.9% favor (*J. oxycedrus*, (*C. funebris, X. vietnamensis*)). This discordance echoes phylogenetic patterns documented in the *Cupressus–Juniperus– Xanthocyparis* complex, where conventional phylogenomic analyses can yield robust but contradictory plastome-based topologies. Zhu et al. (2018) suggested that robust conflicts among plastome-based phylogenies reflect ancient introgression and hypothesized that such events were facilitated by recombination between divergent plastid haplotypes within ancestral heteroplasmic lineages. This process may have generated persistent phylogenetic signals that remain difficult to reconcile with incomplete lineage sorting alone. This small cactus is subsequently merged into a reticulated cherry with *Platycladus orientalis* (Oriental arborvitae), with w_1_=0.126. Finally, the *Thuja*/*Thujopsis* clade, *Fokienia hodginsii* and the clade comprising all other ingroups are grouped into a basal cactus structure. The reference species tree supports *F. hodginsii* and *P. orientalis* as sister lineages; however, our multilocus statistics indicate that the local signal is heterogeneous across loci, with substantial support for competing splits. Specifically, considering the three-taxon set (*C. funebris, F. hodginsii*, and *Thujopsis dolabrata*), 978 loci support the split (*C. funebris, F. hodginsii*) | *T. dolabrata*, whereas 498 loci support (*F. hodginsii, T. dolabrata*) | *C. funebris* (with the remaining 296 loci supporting (*C. funebris, T. dolabrata*) | *F. hodginsii*). The predicted reticulation resulting from this asymmetry between the two minor triplets has been hypothesized by Liu et al. (2022) who constructed a SuperQ (Grünewald et al. 2013) network on a related dataset. A maximum-likelihood tree inferred from the concatenated alignment with IQ-TREE 2 shows that the branches leading to *F. hodginsii* and the (Fig. 7). This raises the suspicion that many of the 498 gene trees supporting(*F. hodginsii, T. dolabrata*) branch leading to *P. orientalis* is significantly longer Thuja lineages are of similar length, while the |*C. funebris* triplet are wrong due to long-branch attraction artefacts. Anyway, TriMouNet retains both dominant resolutions as a reticulation signal.

**FIG. 7.**
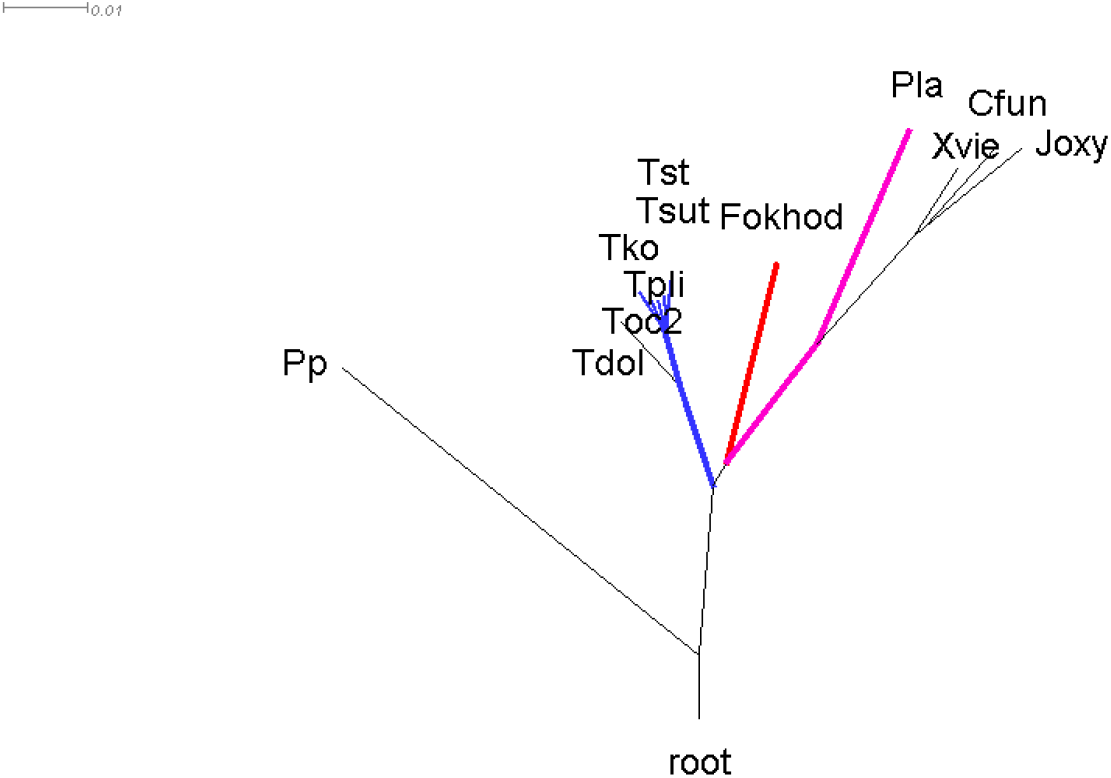
SplitsTree visualization of the IQ-TREE 2 concatenation phylogeny from single-copy nuclear genes, highlighting comparable terminal branch lengths for *Fokienia hodginsii* (red) and the *Thuja* clade (blue).

#### TriMouNet analysis of bird data

We analyzed a phylogenomic dataset comprising 3,679 ultraconserved element (UCE) loci from 11 modern birds (Neoaves) taxa and chicken as outgroup (Jarvis et al. 2014). The neoaves phylogeny is highly controversial (see Stiller et al. (2024) for a recent prominent publication). All taxa included in our networks belong to the generally accepted Telluraves (landbirds) clade. There are seven clusters in the taxa set that have been consistently recovered throughout all recent major studies: parrots (Budgerigar and Kea), parrots+rifleman (Psittacopasseres), Psittacopasseres+falcon (Eufalconimorphae), Eufalconimorphae+Seriema (Australaves), eagle+vulture (Accipitrimorphae), Cuckoo Roller+Trogon (Cavitaves), and the complement of the Australaves (Afroaves). TriLoNet (Fig. 8(i)) failed to resolve any internal phylogenetic split—collapsing all species into a single, unresolved cactus structure. This reflect that for this dataset, with very short interior edges and high variance of the root-to-tip distances, The trinet assignment of TriLoNet is not appropriate. TriMouNet (Fig. 8(ii)) successfully recovered all clear clusters mentioned above and in addition groups

**FIG. 8.**
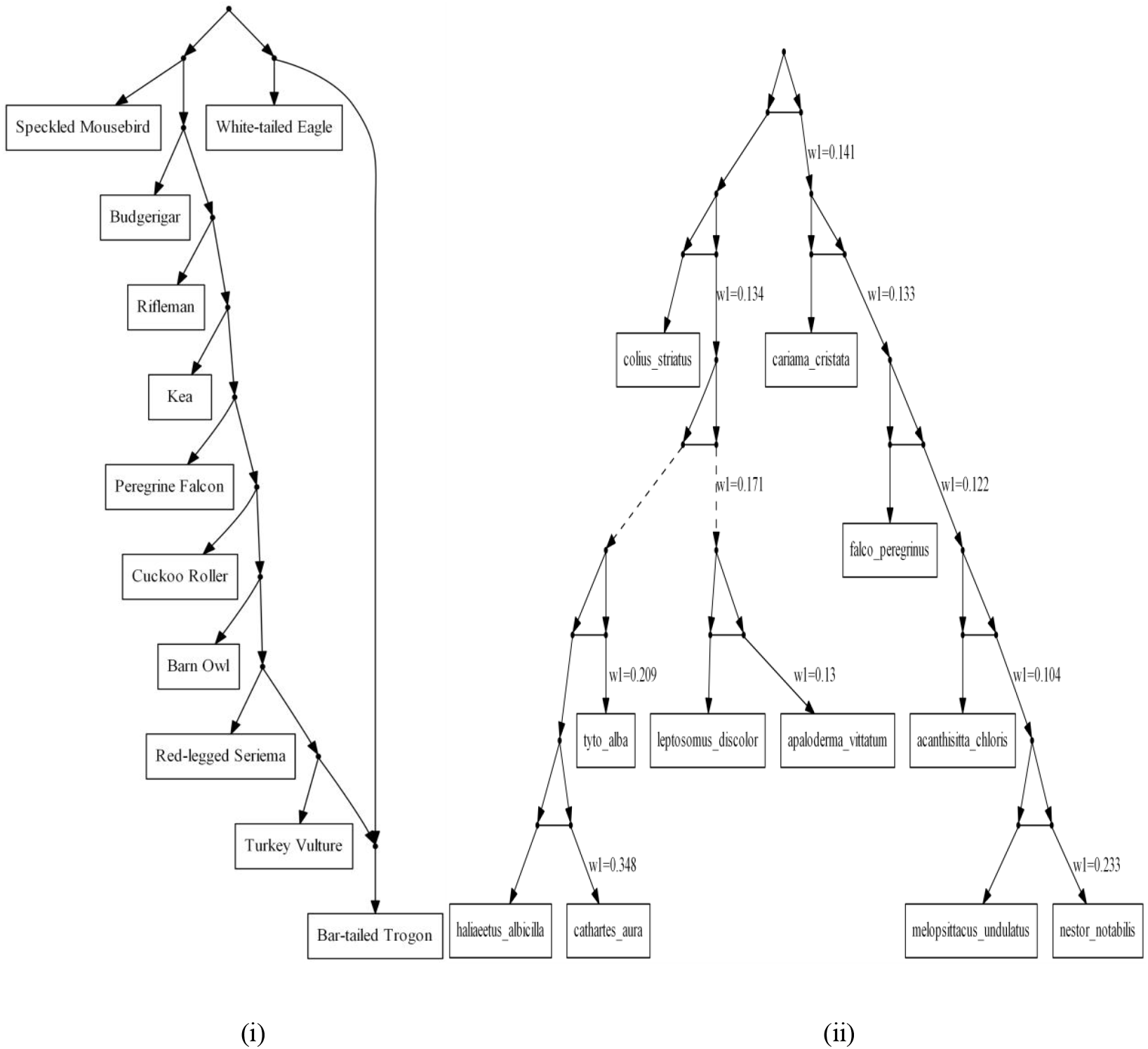
Phylogenetic network of birds based on 3679 Ultraconserved Element genes inferred by TriLoNet and TriMouNet.

Accipitrimorphae with owls and clusters all Afroaves taxa other than mousebird. The former cluster is included in the main tree by Stiller et al. (2024), and the latter one in the UCE tree of Jarvis et al. (2014). Notably, all bifurcations are represented as reticulated cherries, and all corresponding w_1_ values are clearly below 0.5, indicating that the signal places greater weight on the higher-level last common ancestor rather than the lower-level coalescent configuration. We interpret these reticulations as a result of continuous admixture in the early stages of the speciation process.

Unfortunately, TriMouNet struggles to detect the Cavitaves cluster. Indeed, it is visualized by a dashed incoming edge, as well as the edge into the (Accipitrimorphae+owls) cluster. Dashed lines arise, if TriMouNet first tries to merge the taxa into a cactus but does not find the support for any cactus ordering (see Methods and Supplementary Materials). Again, the explanation why the Cavitaves cluster is originally missed, appears to be long-branch attraction because the Cuckoo Roller has a much shorter distance to the last common ancestor than the Bar-tailed Trogon or the Barn Owl.

## Conclusion

We present and implement a new method for phylogenetic network inference, TriMouNet. TriMouNet is built on the general workflow of TriLoNet: it first infers trinets on all three-taxon subsets and then merges them into a network on all taxa, aiming to reconstruct level-1 network structure. The main difference is how trinets are obtained. TriLo-Net assigns trinets directly from three-way sequence alignments, whereas TriMouNet infers gene trees from multi-locus data and then uses the observed distributions of splits and branch lengths to classify and assign trinets. In addition, when we infer gene trees for each three-taxon set, we always include an outgroup. This four-taxon setting helps anchor the topology and, in practice, reduces mistakes that can occur when only three taxa are used. This is important because three-way alignments have limited information: they contain no parsimony-informative patterns, the number of singleton sites strongly depends on branch lengths, and even small departures from a molecular clock can make it hard to identify the strongest triplet, or even the correct network, which can lead to wrong trinet assignments.

In simulations, TriMouNet showed the expected performance across the three signal-strength settings. We also applied TriMouNet to three empirical datasets and compared the results with TriLoNet. Overall, the networks inferred by TriMouNet agree better with known (or better supported) evolutionary relationships and show fewer clear topological errors. That said, TriMouNet can still be affected by errors in the input gene trees. For some loci, compositional heterogeneity, model mismatch, or long-branch attraction can produce incorrect gene trees, and these errors can influence the merged network. Even with this limitation, TriMouNet shows a clear improvement in accuracy and robustness compared with Tri-LoNet.

Future work should focus on improving the inference and quality control of gene trees, for example by using better models and more reliable gene-tree methods together with simple, automated filters to remove low-quality loci, extreme branchlength outliers, and highly inconsistent gene trees; this should further improve the accuracy and stability of trinet inference and network merging.

In a post-processing step, it would be useful to assign trees that are embedded in the TriMouNet to the genes. If it is true that all genes evolved along a single tree and the reconstructed network is correct and complete, then a tree within the network should fit any gene alignment almost as good as the input gene tree. This might help to distinguish correct reticulations from false positives and possibly identify the kind of reticulation that occurred.

Future development of TriMouNet could explore its extension to level-2 networks, a direction supported by recent theoretical and algorithmic results. For example, polynomial-time algorithms for reconstructing rooted binary level-2 networks from trinets have been established for the case where the full set of induced trinets is available (van Iersel et al. 2022). In parallel, work on four-taxon subnetworks shows that semi-directed binary level-2 networks are encoded by their induced quarnets, suggesting that higher-order local structures may help in resolving more complex reticulation patterns (Huber et al. 2025). Taken together, these advances suggest that targeting level-2 network classes is increasingly feasible, although adapting such frameworks to noise and missing information in genomic-scale data—while maintaining TriMouNet’s computational efficiency—remains a key challenge.

## Materials and Methods

TriMouNet modifies the trinet of TriLoNet to match what is identifiable from multilocus dataset. In the S_1_ trinet, the branch-length and split distributions allow us to determine which taxon is below the reticulation node, so we retain a directed reticulation structure. In contrast, for the S_2_ trinet, these distributions do not resolve which taxon is below the reticulation node, and we therefore connect the two relevant internal nodes by an undirected horizontal edge. Similarly, for NT-type trinets, the branch-length and split distributions do not identify the relative hierarchical ordering of the corresponding internal nodes, which are the last common ancestors of the involved lineages; accordingly, we represent their relationship with an undirected horizontal edge between the two internal nodes.

As a result, TriLoNet’s N_1_ and N_2_ are grouped as our N_2_, TriLoNet’s N_4_ and N_5_ are grouped as our N_4_, while TriLoNet’s N_3_ and T_1_ correspond to our N_3_ and T_1_, respectively. In TriMouNet, binets are classified according to the branch-length distributions between two taxa, with S_0_ and T_0_ corresponding to bimodal and unimodal distributions, respectively. Trinets are classified based on branch-length and split distributions among three taxa, where NT-type trinets display a single dominant split and S-type trinets display two competing splits. Altogether, this leads to six distinct trinet types and two binet types in our framework (Fig. 1).

### Pending Edge, Cherry Edge and Common Edge

Before introducing the binet and trinet types, we define three branch-length summaries that will be used throughout our inference of branch-length distribution patterns.

For an NT-type trinet, the relevant summaries are the cherry edge and the pending edge. Specifically, for a triple {x, y, z}, let the most supported split be xy|z. We define the cherry edge as the distance from x or y to the most recent common ancestor of x and y. We define the pending edge as the distance from z to the most recent common ancestor of x, y and z. Using Fig. 9 as an illustration, the N_2_ (x, y; z) has a cherry edge of a, while the pending edge can take values a+b or a+b+c depending on the displayed tree. For N_3_ (x; y, z), the cherry edge can be a or a+b, and the pending edge is a+b+c.

**FIG. 9.**
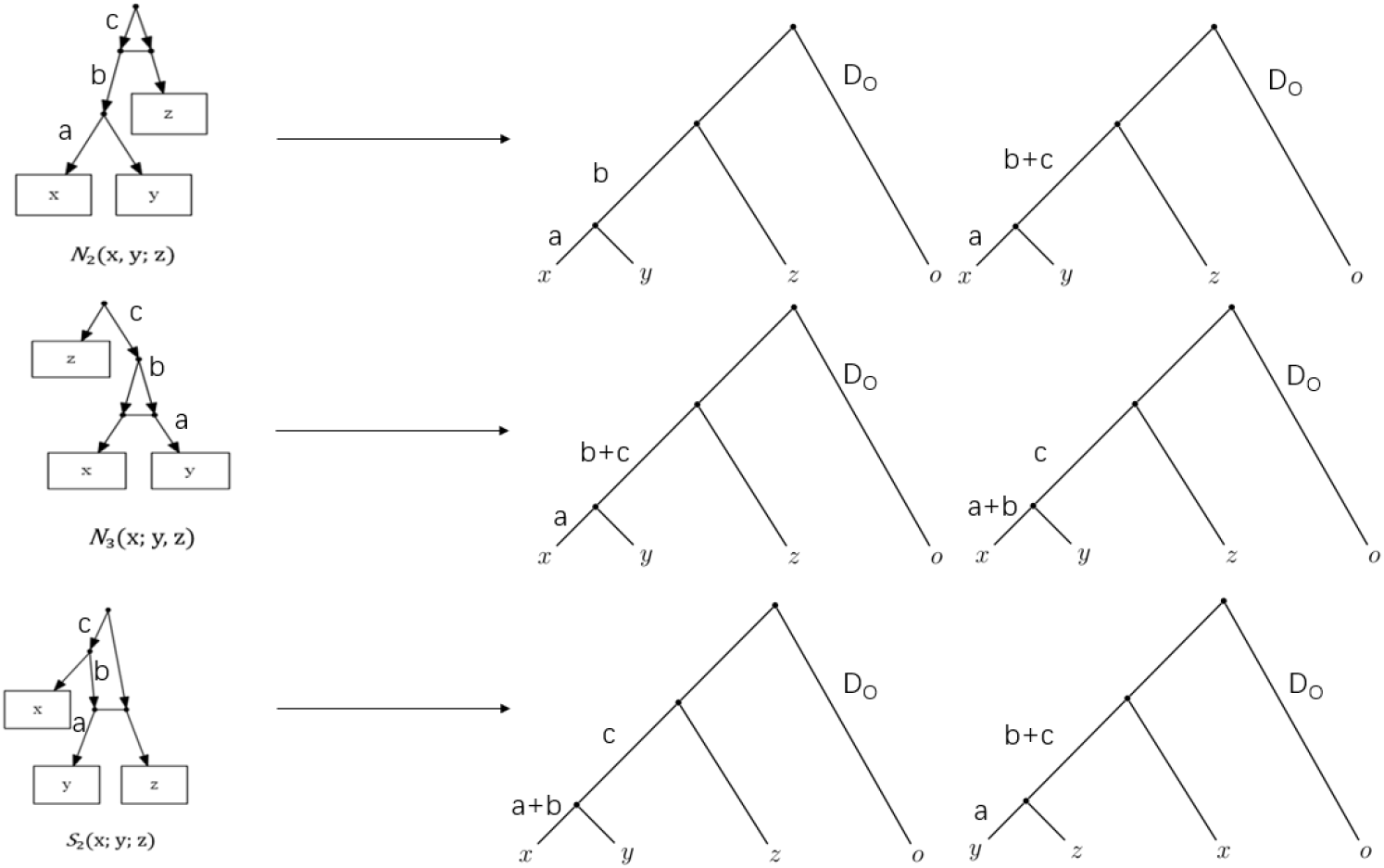
Decomposition of N_2_, N_3_ and S_2_ based on the horizontal reticulation edge.

For S-type trinet, we instead use the common edge, which is defined when two alternative splits are supported. For example, if the two most supported splits are xy|z and yz|x, then y is the taxon shared by the two alternative cherries (x, y) and (y, z). We define the common edge as the distance from this shared taxon y to its parent in the corresponding displayed trees. Using Figure 9. as an illustration, the common edge equals a or a+b.

### Binet and Trinet Types

#### Binet Structures (S_0_ and T_0_)

In our framework, the S_0_ binet is redefined by placing the two sub-ancestors of taxa x and y at the same hierarchical level, connected by a single undirected horizontal edge, with the higher ancestor linking independently to both, yielding a mixture branch length distribution. If the two sub-ancestors coincide, the structure reduces to a T_0_ binet, corresponding to a single branch length distribution.

#### Trinet Structures (NT-type trinet)

In all four trinet types, sister taxa x and y first coalesce through a sub-ancestor, followed by coalescence with taxon z at a higher ancestor. Conceptually, an NT-type trinet can be viewed as the combination of two binets associated with the same sister-taxon pair. Different combinations of these two binet types give rise to four subclasses of NT-type trinets, denoted as N_2_, N_3_, N_4_, and T_1_. The four types are described as follows:

T_1_: Sister taxa x and y are connected via a T_0_-type binet, and their ancestor then connects to taxon z with the other T_0_-type binet, producing single branch length distributions for both cherry edge and pending edge.

N_2_: Sister taxa x and y are connected via a T_0_-type binet, and their ancestor is then connected to taxon z with the other S_0_-type binet, producing a single branch length distribution for the cherry edge and a mixture branch length distribution for the pending edge.

N_3_: Sister taxa x and y are connected via a S_0_-type binet, and their ancestor is then connected to taxon z with the other T_0_-type binet, producing a mixture branch length distribution for the cherry edge and a single branch length distribution for the pending edge.

N_4_: Sister taxa x and y are connected via a S_0_-type binet, and their ancestor then connects to taxon z with the other S_0_-type binet, producing mixture branch length distributions for both cherry edge and pending edge.

#### Trinet Structures (S-type trinet)

S-type trinets capture more structured and asymmetric evolutionary relationships and are characterized by displaying two splits. Two subclasses of S-type trinets are distinguished in our framework, denoted as S_1_ and S_2_, which differ in whether the last common ancestors of the two alternative cherries occur at the same hierarchical level or at different levels in the underlying network.

The two types are described as follows:

S_1_: The first two coalescent sub-ancestors are at the same hierarchical level, producing a single branch length distribution for the common edge (z) shared by the two cherry groups: (x, z) and (y, z).

S_2_: The first two coalescent sub-ancestors are at different hierarchical levels, resulting in two different branch length distributions for the common edge (y) shared by the two cherry groups: (x, y) and (y, z).

### Branch-length distribution assumption and EMG likelihood

We model the distribution of estimated gene-tree branch lengths using an exponentially modified Gaussian (EMG) distribution (Grushka et al. 1972). Under the multispecies coalescent, the coalescent time T on an internal branch follows an exponential distribution with rate λ and mean 1/λ=2N_e_ (Hudson 1983; Tajima 1983), where N_e_ is the effective population size. In empirical analyses, the observed branch length also contains estimation error due to finite sequence length, model misspecification, all of which contribute to stochastic deviations from the true value (Hibbins et al. 2023). Because branch-length estimation aggregates information over many approximately independent sites, the cumulative estimation error E can be approximated as Gaussian, E∼N(μ, σ^2^). Thus, the observed branch length L is modeled as L=T+E, where T∼Exp(λ) and E∼N(μ, σ^2^). The resulting distribution of L is the exponentially modified Gaussian distribution, L∼EMG(u, σ, λ). where μ represents the mean of the Gaussian component, σ is the standard deviation of the estimation error, and λ controls the exponential decay associated with coalescent waiting times.

### Normalized Distance Variables (V_1_, V_2_ and V_3_)

To capture the coalescent patterns of sister species within a quartet, we defined three normalized distance variables, V_1_, V_2_ and V_3_ derived from the reconstructed gene tree branch lengths.

V_1_ quantifies the relative tree distance between the sister species (x, y) within the 3-taxa tree induced by x, y and the outgroup o:

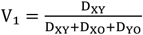

where D_XY_ is the sum of the cherry edges between x and y, and D_XO_, D_YO_ are the branch lengths from taxon x and y to the outgroup taxon O, respectively.

V_2_ captures the relative distances between one member of the sister pair and a third species z, averaged across the two possible pairings:

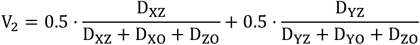

where D_XZ_ and D_YZ_ are the path lengths from taxon x or y to taxon z. V_2_ therefore represents an averaged normalized distance from the sister pair to the third species in the quartet.

V_3_ represents the ratio of the length of the common edge (D_C_) to the length of the path from the common taxon C to the outgroup o, which includes the length of the interior edge (D_I_) and the length of the pending edge incident with the outgroup o (D_O_).

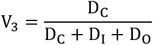

We model the estimated branch length L as L=T+E, L follows an exponentially modified Gaussian distribution. The normalized variables V_1_, V_2_, V_3_ are used as observations of L for parameter inference under the EMG model.

By fitting an EMG to these normalized variables across multiple gene trees, we can statistically evaluate whether the observed patterns are consistent with a single or a mixture of two coalescent time scenarios.

### Assign Trinets from multi-locus gene trees (Step A)

Model-based multi-locus gene trees derived trinets offer greater accuracy and biological interpretability than sequence pattern based trinets, which are constrained by limited site information and substantial stochastic noise. In our approach, we apply a model-based method to reconstruct multilocus gene trees from a taxon set X of size n that includes an outgroup o, which could be considered an extension of TriLoNet.

#### Step A1

In every reconstructed gene tree, we obtain (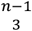) quartet trees by taking the induced subtree on {x,y,z,o} from the full gene tree and contracting redundant internal edges. For each triple {x, y, z}, we denote s(xy|z) as the number of gene trees supporting topology ((x,y),z,o), which corresponds to the split xy | z; s(xz|y) for topology ((x,z),y,o), which corresponds to the split xz | y; s(yz|x) is for topology ((y,z),x,o), which corresponds to the split yz|x. Without loss of generality, we order them such that s(xy|z) ≥ s(xz|y) ≥ s(yz|x).

We perform a binomial test with null hypothesis H_0_: p=0.5 by comparing the second-highest count s(xz|y) against s(xz|y)+s(yz|x), and denote the resulting *p*-value by P_s_ .

The value of P_s_ is used to determine whether the triple is assigned to 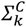 (S-type trinets) or to ∑_*k*_ (NT-type trinet). A triple is assigned to ∑_*k*_ if P_s_≥δ_1_, indicating that the observed asymmetry can be explained by chance under H_0_. If P_s_ < δ_1_ the asymmetry between these two counts is too large to be explained by random sampling, which is characteristic of S-type trinets, the corresponding triple is therefore assigned to 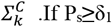, the two counts do not differ significantly, indicating a nearly symmetric signal typical for NT-type trinets, and the triple is assigned to ∑_*k*_.

To control the family-wise Type I error arising from performing (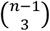) binomial tests, we apply a Bonferroni correction and set the significance threshold to δ_1_=0.05/^(*n*−1)^.

For completeness, we also define P_f_ analogously by comparing the highest count s(xy|z) against s(xy|z)+s(xz|y); P_f_ is not used to assign trinets but will be employed in a subsequent step of the algorithm.

#### Step A2

In this step, we need to compute the two normalized distance variables V_1_ and V_2_ in all triples which belong to ∑_*k*_. We then compute the probability density function of the exponentially modified Gaussian distribution for each gene tree based on these data, take the negative logarithm, and sum them across loci. We will apply one μ(μ_1_) or two μ(μ_1_ and μ_2_) to construct the objective function of ∑_*k*_. In order to reduce the number of free parameters, we assumed that the variance and the λ parameters are equal for both components of the mixtures.

##### Probability density function

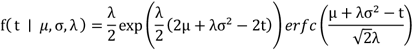

##### Objective function of ∑_*k*_

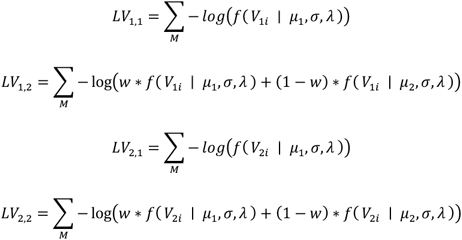

##### Objective function of 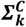

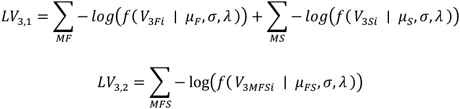

In the objective function, we have the values of V_1_ and V_2_ for all genes as the input and 0<w<1 as the weight of μ_1_. V_3_ is the observed dataset in common edge of type S. M means the number of loci in each of triple. MF means the number of loci supports first split xy|z. MS means the number of loci supports second split. MFS means the number of loci supports first or second triple.

#### Step A3

In this step, we use the constrained L-BFGS-B algorithm (Broyden et al. 1970) to obtain the optimized objective value and the corresponding parameters. In practice, when fitting the two-mean EMG mixture using L-BFGS-B, the parameters σ and λ are often strongly coupled with the means u_1_ and u_2_. Under the constrained optimization setting, this coupling frequently prevents the algorithm from reaching a global optimum. To address this issue, we transform the variables into the objective function and replace the original parameters with unconstrained variables. The detailed reparameterization procedure follows the standard formulation of mixture distributions and is provided in the Supplementary Materials.

#### Step A4

We need to assign a trinet to each triple t = {x, y, z} in ∑_*k*_ according to the gap of objective function value between one μ(μ_1_) or two μ (μ_1_ and μ_2_) with V_1_ and V_2_. If |Lv_1,1_-Lv_1,2_| ≤ κ, it means the relative tree distance between the sister species (x, y) is unimodal distribution, which corresponds to T_1_ (x, y, z) or N_2_ (x, y; z), otherwise is N_3_ (x; y, z) or N_4_ (x; y; z). If |Lv_2,1_-Lv_2,2_ | ≤ κ, it means the relative distances between one member of the sister pair and a third species is unimodal distribution, which corresponds to T_1_ (x, y, z) or N_3_ (x; y, z), otherwise is N_2_ (x, y; z) or N_4_ (x; y; z). According to |Lv_1,1_-Lv_1,2_| and |Lv_2,1_-Lv_2,2_ |, we could determine the type of each triple.

#### Step A5

We also need to assign a trinet to each triple t = {x, y, z} in 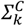 according to objective function value between the sum of separated two groups and the combined group with V_3_. If |Lv_3,1_-Lv_3,2_|≤ κ, it means the common edge between the first sister species (x, y) and the second sister species (x, z) is unimodal distribution, which corresponds to S_1_ (x, y; z), otherwise is S_2_ (x; z; y).

### Merge Trinets to construct a level-1 phylogenetic network (Step B)

Similar to TriLoNet, our method uses Tarjan’s algorithm (Tarjan 1972) to compute the strongly connected components of the graph, thereby identifying a non-singleton subset Y of X that corresponds to a cherry, reticulated cherry or cactus. In addition, we incorporate parameters estimated with the L-BFGS-B algorithm. We process each identified subset Y iteratively and assemble a level-1 phylogenetic network, using the associated trinet information to guide the construction.

#### Step B1

At the start of the algorithm, we input the trinets obtained from Step A. Each trinet records both its inferred type and the L-BFGS-B -optimized parameters estimated in Step A. We construct a matrix M that reflects the degree of association between species by using the dense collection of trinets T and X. Specifically, for each trinet T_i_∈T defined on the taxon set {x_i_,x_j_,x_k_}⊆X, we update the association matrix M=(m_ab_)∈R^|X|×|X|^ according to the inferred trinet type. Based on the inferred trinet type and the corresponding P_f_ values estimated in Step A, we define a new rule to construct the association matrix M. This rule integrates both the categorical information of trinet types and the quantitative support provided by the estimated parameters. We then construct the directed graph D by applying a threshold δ_2_=|X|-2+log(P_B_) to the entries of M, where 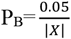. This choice of δ_2_ follows a statistical principle for controlling the Type II error rate. Finally, the identification of a non-singleton subset Y follows the same framework as in TriLoNet. Full details of identifying non-singleton subsets Y of X are provided in the Supplementary Materials.

#### Step B2

We need to construct a network N_Y_ from subset Y in step B1. If |Y|=2, let Y={x, y}, N_Y_ will either be a binet S_0_(x; y) or a binet T_0_(x, y). For each trinet T_i_ in which taxa x and y form a sister pair, we use the average of |Lv_1,1_-Lv_1,2_| to determine whether N_Y_ is reticulated or not. If |Y|≥3, N_Y_ is constructed either as a cactus or as a collection of binets, depending on the types of the associated trinets T_i_ induced by triples {x_i_, y_i_, z_i_} ⊆ Y. In particular, we build N_Y_ as a cactus when there are enough S-type trinets so that every taxon in Y appears in at least one S-type trinet. We then apply ordering rules so that the two sides of the cactus and the bottom leaf agree, as much as possible, with the trinet information associated with Y. Full details of the binet and cactus construction are given in the Supplementary Materials. Otherwise, we restrict the construction by enforcing a stricter criterion in Step B1, setting the threshold δ_2_ from |X|-2+log(P_B_) to |X|-2, which yields a subset Y^∗^⊂Y. We then apply the same construction strategy to obtain the induced subnetwork N_Y*_. When the available information does not support a cactus, we display the corresponding part of the inferred network using dashed edges instead of solid ones.

#### Step B3

We will merge every taxon in Y into a new taxon y^*^, obtain the new taxa set X^*^ and the trinet set T^***\****^ induced by X^*^, and then use X* and T* to continue constructing the network N^*^. We then combine N_Y_ and N^*^, repeat Steps B1–B2 recursively until the identified non-singleton subset satisfies Y=X.

## Implementation

TriMouNet (TriMouNet.jar) is freely available at https://github.com/Ronin-Mqy/TriMouNet. This repository also contains the simulation datasets and the empirical data used in this study. The software requires users to first infer multilocus gene trees and merge them into a single tree file as input. A provided Python pipeline then processes the gene-tree collection and outputs a .tnet file, which records the trinets assigned in Step A as well as the associated parameter estimates required in Step B. Given this .tnet file, TriMouNet generates a network in DOT format, which can be visualized using Dendroscope (Huson and Scornavacca 2012) and GraphViz (Ellson et al. 2002), respectively.

## Supplementary Material

Supplementary material is available as a separate PDF file.

