## Supplementary Material for "TriMouNet: An Algorithm for Inferring Level-1 Phylogenetic Networks from Multi-Locus Gene Tree Distributions"

In this Supplementary Material, we divide the content into three parts.

The first part describes the variable transformation used in Step A3. In the original formulation, the four parameters $\mu_1$, $\mu_2$, $\sigma$, and $\lambda$ are not mutually independent. When $\mu_1$ and $\mu_2$ are optimized simultaneously, this dependency restricts the feasible optimization range of $\sigma$ and $\lambda$, which may prevent the L-BFGS-B algorithm from reaching the global optimum. To address this issue, we transform these four dependent parameters into three independent variables: x, y, and w, thereby improving the stability and effectiveness of the L-BFGS-B optimization and facilitating convergence to the global optimum.

The second part provides technical details of the merging procedure. Specifically, we describe how matrix representations are used to identify non-singleton subsets Y, how the intermediate network $N_Y$ is constructed when $N_Y$ corresponds to a cherry, a reticulated cherry, or a cactus, and how the associated trinets are reduced during this merging process.

The third part is a threshold experiment that evaluates a simple locus-number–dependent cutoff for distinguishing single and mixture signals in trinet inference.

### Variable reduction and transformation

Let $X_1$ and $X_2$ be independent random variables following exponentially modified Gaussian distribution (EMG, also known as exGaussian distribution). $X_1 \sim EMG(\mu_1, \sigma_1^2, \lambda_1)$ and $X_2 \sim EMG(\mu_2, \sigma_2^2, \lambda_2)$.

Therefore, a mixture $wX_1+(1-w)X_2$ has seven free parameters, which are difficult to optimize in practice. We reduce the dimension of the problem to three by assuming $\lambda:=\lambda_1=\lambda_2$, $\sigma:=\sigma_1=\sigma_2$, and by assuming that the mean and the variance of the mixture equals the sample mean and the sample variance, respectively. In addition, we transform the remaining variables to simplify the search spaces.

Let X denote a random variable drawn from a two-component mixture of $X_1$ and $X_2$. Let $S_m$ and $S_v$ denote the mean and variance of X, respectively, while $S_{m1}$, $S_{m2}$; $S_{v1}$, $S_{v2}$ and $w_1$, $w_2$ denote the means, the variances and the weights of the first component and the second component;

In the exGaussian model, $\mu$ denotes the mean of Gaussian component, $\sigma^2$ denotes the variance of Gaussian component, $\lambda$ denotes the rate parameter of the exponential component. We reparameterize the model by defining $x=1/\lambda$, which is used as the optimized parameter. Since the expectation of an EMG equals $\mu+1/\lambda$, we have

$E(X)=S_m= w_1S_{m1}+ w_2S_{m2}=w_1(\mu_1+x)+w_2(\mu_2+x)=w_1\mu_1+w_2\mu_2+x.$

Since the variance of an EMG equals $\sigma^2+1/\lambda^2$, the variance D(X) of X equals

$D(X)=S_v=E(X^2)-S_m^2=w_1(S_{v1}+S_{m_1}^2)+ w_2(S_{v2}+S_{m_2}^2)-S_m^2$

$=w_1(\sigma_1^2+x^2+(\mu_1+x)^2)+ w_2(\sigma_2^2+x^2+(\mu_2+x)^2)-S_m^2$

$= w_1(\sigma_1^2+\mu_1^2)+ w_2(\sigma_2^2+\mu_2^2)+2S_mx-S_m^2$

This implies $S_v+S_m^2= w_1(\sigma_1^2+\mu_1^2)+ w_2(\sigma_2^2+\mu_2^2)+2S_mx$ (1)

To enforce the linear constraint $w_1\mu_1+w_2\mu_2=C=S_m-x$, we introduce a one-dimensional parameterization $\mu_1=C-w_2t$, $\mu_2=C+w_1t$ (2), which satisfies the constraint for all $t\in R$.

Substituting (2) into (1), we obtain

$w_1\sigma_1^2 + w_2\sigma_2^2+C^2+2w_1w_2t^2= S_v+S_m^2-2S_mx$ (3)

From (3), we have $S_v+S_m^2-2S_m x \geq (S_m-x)^2=C^2$, thus $x \leq \sqrt{s_v}$

Moreover, the non-negativity of the left-hand side of (3) implies $S_v+S_m^2-2S_m x \geq 0$, which yields $x \leq \frac{s_v}{2s_m}+\frac{s_m}{2}$

Combining this result with the previous bound, we obtain

$x \leq \min(\sqrt{s_v}, \frac{s_v}{2s_m}+\frac{s_m}{2})$

From (1) and (3), we get $w_1\mu_1^2+ w_2\mu_2^2= C^2+ w_1w_2t^2$ (4)

Let $y^*=\frac{w_1\sigma_1^2+w_2\sigma_2^2}{S_v+S_m^2-2S_m x}$, so $t^2=\frac{(S_v+S_m^2-2S_m x)(1-y^*)-C^2}{w_1 w_2}$, (5)

Since $w_1+w_2=1$, the AM–GM inequality implies that $w_1w_2 \leq 1/4$

Consequently, $t \geq 2\sqrt{(S_v + S_m^2 - 2S_m x)(1 - y^*) - C^2}$ (6)

From (5), we get $(S_v + S_m^2 - 2S_m x)(1 - y^*) - C^2 \geq 0 \Rightarrow y^* \leq \frac{S_v-x^2}{(S_v+S_m^2-2S_m x)}$

Let $y= (\frac{S_v+S_m^2-2S_m x}{S_v-x^2})y^*$, so $y \leq 1$

From (2) and (5), we could get $\mu_1= S_m-x-\sqrt{\frac{((S_v+S_m^2-2S_m x)(1-y^*)-C^2)w_2}{w_1}}$ (7)

From (7) we could get $S_m-x \geq \sqrt{\frac{((S_v+S_m^2-2S_m x)(1-y^*)-C^2)w_2}{w_1}} \Rightarrow \frac{((S_v+S_m^2-2S_m x)(1-y^*))w_2}{(Sm-x)2} \leq 1$

Let $w=\frac{((S_v+S_m^2-2S_m x)(1-y^*))w_2}{(Sm-x)2}$, $0<w \leq 1$

$\Rightarrow w_1=1-\frac{w(Sm-x)2}{(S_v+S_m^2-2S_m x)(1-y^*)} = 1-\frac{w(Sm-x)2}{S_v+S_m^2-2S_m x-S_v y+x^2 y}$

$w_2=\frac{w\,(Sm-x)2}{((S_v+S_m^2-2S_m x)(1-y^*))} = \frac{w\,(Sm-x)2}{S_v+S_m^2-2S_m x-S_v y+x^2 y}$(8)

From (5), we get

$\sigma_1=\sigma_2=\sqrt{y^*(S_v + S_m^2 - 2S_m x)}=\sqrt{y(S_v - x^2)}$

From (5) and (8), we get

$\mu_1=C-w_2t=C-\frac{w(Sm-x)2}{S_v+S_m^2-2S_m x-S_v y+x^2 y}\sqrt{\frac{S_v-x^2-S_v y+x^2 y}{(1-\frac{w(Sm-x)2}{S_v+S_m^2-S_m x-S_v y+x^2 y})\frac{w(Sm-x)2}{S_v+S_m^2-S_m x-S_v y+x^2 y}}}$,

$\mu_2=C+w_1t= C+(1-\frac{w(Sm-x)2}{S_v+S_m^2-2S_m x-S_v y+x^2 y})\sqrt{\frac{S_v-x^2-S_v y+x^2 y}{(1-\frac{w(Sm-x)2}{S_v+S_m^2-2S_m x-S_v y+x^2 y})\frac{w(Sm-x)2}{S_v+S_m^2-2S_m x-S_v y+x^2 y)}}}$

After this reparameterization, we obtain three independent variables x, y, and w which can be optimized without constraints among them. We apply the L-BFGS-B algorithm in this parameter space. After this transformation, the optimization procedure achieves stable convergence and reduces the risk of convergence to local optima or spurious solutions.

### Detecting Small sink sets

Given a collection of trinets T defined on the taxon set X, this procedure identifies a non-singleton subset $Y \subseteq X$ corresponding to a small SN-set. We compute a matrix $M = M_{ij}$ that assigns a non-negative association value to every pair (i, j) of taxa. M is initialized as an all-zero matrix.

If $T_i$ corresponds to an $S_1$ or $S_2$ trinet on $\{x_i,x_j,x_k\}$, all pairwise associations among the three taxa are strengthened by one, that is,

$$\begin{pmatrix} m_{ii} & m_{ij} & m_{ik} \\ m_{ji} & m_{jj} & m_{jk} \\ m_{ki} & m_{kj} & m_{kk} \end{pmatrix} \longleftarrow \begin{pmatrix} m_{ii}+1 & m_{ij}+1 & m_{ik}+1 \\ m_{ji}+1 & m_{jj}+1 & m_{jk}+1 \\ m_{ki}+1 & m_{kj}+1 & m_{kk}+1 \end{pmatrix}.$$

If $T_i$ corresponds to an NT-type trinet, we use a $P_f$ value inferred by Step A to represent the degree of association between $x_k$ and $x_i$ or $x_j$. In this case, the associations from i and j to k are reduced by $-\log(P_f)$, while all other association within i, j, k are increased by one.

$$\begin{pmatrix} m_{ii} & m_{ij} & m_{ik} \\ m_{ji} & m_{jj} & m_{jk} \\ m_{ki} & m_{kj} & m_{kk} \end{pmatrix} \longleftarrow \begin{pmatrix} m_{ii}+1 & m_{ij}+1 & m_{ik}+\log(p_f) \\ m_{ji}+1 & m_{jj}+1 & m_{jk}+\log(p_f) \\ m_{ki}+1 & m_{kj}+1 & m_{kk}+1 \end{pmatrix}.$$

After all trinets have been processed, we set all diagonal entries of M to zero, $m_{aa} \leftarrow 0$ for all $a \in X$, thereby removing self-associations and retaining only interspecific relationships for subsequent steps of the algorithm.

A naive thresholding rule that compares each entry $m_{ij}$ against the hard cutoff $|X|-2$ is overly conservative. We apply a Bonferroni-style correction and replace the threshold with

$$\delta_2 = |X| - 2 + \log(\frac{0.05}{|X|})$$

which provides an explicit criterion for controlling the Type II error rate under multiple pairwise decisions. If $m_{ij} \geq \delta_2$, we insert a directed edge from taxon i to taxon j, thereby constructing a directed graph D. We then apply Tarjan's algorithm to compute the strongly connected components and extract a non-singleton subset Y. All subsequent steps, including construction of the condensed graph and selection of Y, follow the same framework as in TriLoNet.

### Assigning binets

Given a collection of trinets T defined on the small sink set Y= {x, y} for each trinet $T_i \in T$, the associated parameters $\mu_1$, $\mu_2$, $w_1$, and $|Lv_{1,1}\text{-}Lv_{1,2}|$, this procedure constructs either a binet $S_0$ or a binet $T_0$.

We iterate over all trinets $T_i \in T$. For each trinet that contains both taxa x and y, we first check whether its inferred type belongs to $T_1$, $N_2$, $N_3$, or $N_4$. In addition, we require that x and y do not occupy the third position of the ordered taxon triple defining the trinet, ensuring that x and y are inferred to form the candidate cherry in $T_i$.

All trinets satisfying the above conditions are collected, and let L denote the total number of such supporting trinets. For these trinets, we compute the averaged values of the parameters $\mu_1$, $\mu_2$, and $|Lv_{1,1}\text{-}Lv_{1,2}|$. The weight parameter $w_1$ is aggregated across the L supporting trinets by taking the L-th root of the product of their individual $w_1$ values.

If the averaged value of $|Lv_{1,1}\text{-}Lv_{1,2}|$ is smaller than the threshold κ defined in Step A4, we construct a reticulated binet $S_0(x, y)$; otherwise, a non-reticulated binet $T_0(x, y)$ is constructed.

When an $S_0(x, y)$ binet is constructed, indicating that taxa x and y form a reticulated cherry, we further annotate this binet with the aggregated parameter estimates inferred from the supporting trinets. Specifically, the aggregated weight parameter $w_1$ is assigned to the reticulation associated with $S_0(x, y)$. These parameter values are retained for subsequent steps of network construction.

### Cactus fitting

Before introducing the algorithm for constructing a cactus, we first describe the properties of a cactus. The supplementary material (Oldman et al. 2016) defined the tuple (a1, a2, . . . , ap : b1, b2, . . . , bq : z), or equivalently (b1, b2, . . . , bq : a1, a2, . . . , ap : z), is the cactus as depicted in Fig. 1. This cactus consists of two sets of taxa: a1, a2, . . . , ap ∈ taxa set A and b1, b2, . . . , bq ∈ taxa set B.

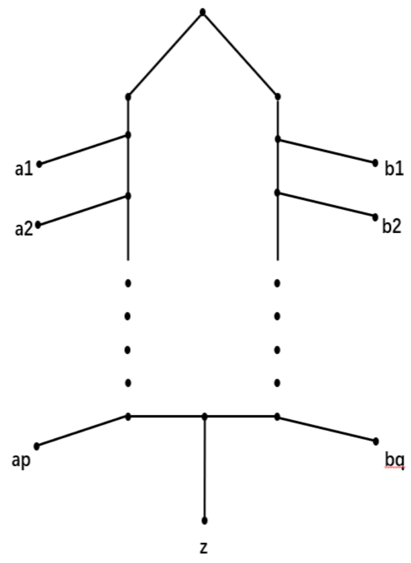


**FIG. 1. Cactus of {a1, . . . , ap, b1, . . . , bq, z}.**

In our algorithm, the objective is to construct a cactus that reflects, to the greatest extent possible, the dense collection of trinets T associated with Y. Specifically, we apply a set of rules to determine the ordering of the two sides of the cactus to maximize the support for $S_2$, the bottom leaf z of the cactus maximum the support for $S_1$ and $S_2$. In each trinet which belongs to $\Sigma_k^C$, if the trinet is $S_1$ (x, y; z), the $S_2$-support score of leaf x and y is $\phi$ (x)= $\phi$ (y)=$|Lv_{3,1}-Lv_{3,2}|-\kappa$, which is lower than 0; if the trinet is $S_2$ (x; y; z), the $S_2$-support score of leaf x and y is $\phi$ (x)= $\phi$ (y)=$|Lv_{3,1}-Lv_{3,2}|-\kappa$, which is higher than 0. The support score for $S_1$ and $S_2$ of leaf z is $\phi$ (z)=-log ($P_s$).

The workflow of our algorithm is as follows:

(i) We let $\phi$ (x)=0 for all x ∈ Y, x is a leaf set from the small sink set Y used in this algorithm. We iterate over all trinets $T_i \in \Sigma_k^C$. For each trinet $T_i$, if $T_i$= $S_1$ (x, y; z) or $T_i$= $S_2$ (x; y; z), we update $\phi$(z) by setting $\phi$ (z)=$\phi$ (z) -log ($P_s$). After processing all such trinets, the taxon g with the maximum value of $\phi$ is selected, and we designate taxon g as the bottom leaf of the cactus.

(ii) We let X = Y - {g}, $\phi$(x)=0 for all x ∈ X. We iterate over all trinets Ti$\in\Sigma_k^C$. For each trinet $T_i$, if $T_i$= $S_1$ (x, y; z) or $T_i$= $S_2$ (x; y; z), we update $\phi$ (x)=$\phi$ (x) + $|Lv_{3,1}-Lv_{3,2}|-\kappa$, $\phi$ (y)=$\phi$ (y) + $|Lv_{3,1}-Lv_{3,2}|-\kappa$. After processing all such trinets, taxon $a_1$ corresponding to the maximum value of $\phi$ is selected, and we designate taxon $a_1$ as the top leaf of left side in this cactus. We assign $a_1$ to taxa set A.

(iii) We let X = X - {$a_1$}, $\phi$(x)=0 for all x ∈ X. We iterate over all trinets $T_i\in\Sigma_k^C$. For each trinet $T_i$, if $T_i$= $S_1$ (x, y; g) or $T_i$= $S_2$ (x; y; g), we update $\phi$ (x)=$\phi$ (x) + $|Lv_{3,1}-Lv_{3,2}|-\kappa$, $\phi$ (y)=$\phi$ (y) + $|Lv_{3,1}-Lv_{3,2}|-\kappa$. After processing all such trinets, taxon x corresponding to the maximum value of $\phi$ is selected, we then need to assign taxon x to taxa set A or B.

(iv) We let X = X - {x}, $\phi$(x)=0 for all x ∈ X. We first iterate taxon $a_j$ from taxa set A, then iterate over all trinets $T_i\in\Sigma_k^C$. For each trinet T, if $T_i$= $S_1$ ($a_j$, x; g) or $T_i$= $S_2$ ($a_j$; x; g) or $T_i$= $S_1$ (x, $a_j$; g), we update $\phi$ (x)=$\phi$ (x) + $|Lv_{3,1}-Lv_{3,2}|-\kappa$. We let $\Phi_A$(x)= $\phi$ (x).

If the size of B is 0, if $\phi$ (x)≤0, we assign x to taxa set B, otherwise, we assign x to taxa set A. If the size of B is not 0, we let $\phi$(x)=0 again. We first iterate taxon $b_k$ from taxa set B, then iterate over all trinets $T_i\in\Sigma_{kc}$. For each trinet T, if $T_i$= $S_1$ ($b_k$, x; g) or $T_i$= $S_2$ ($b_k$; x; g) or $T_i$= $S_1$ (x, $b_k$; g), we update $\phi$ (x)=$\phi$ (x) + $|Lv_{3,1}-Lv_{3,2}|-\kappa$. We let $\Phi_B$(x)= $\phi$ (x). If $\Phi_A(x)\geq\Phi_B(x)$, we assign taxon x to taxa set A, else, we assign taxon x to taxa set B.

### Trinet merging

After we construct a network $N_Y$ by assigning a binet or fitting a cactus, we perform a trinet-merging step to aggregate the associated trinets into a single modified trinet that best represents the underlying signal distribution inferred from the estimated trinet parameters.

In this step, a key task is to determine both the type and the relative order of each merged trinet. Specifically, let Y= $\{y_1, y_2, \ldots, y_m\}$ denote the set of taxa to be merged, and let X'= $\{x_1, x_2, \ldots, x_n\}$ denote the remaining taxa. We consider all triples of the form t'= $\{x_i, x_j, y_k\}$ which contains exactly one taxon from Y and two taxa from X'. For a fixed pair $\{x_i, x_j\}$, the m such triples obtained by varying $y_k \in Y$ are merged into a single representative trinet. To determine the type of the merged trinet, we aggregate the statistical evidence across m constituent triples by computing

$$P^* = \sqrt[m]{\prod_{k=1}^{m} \mathrm{P}_\mathrm{s}^\mathrm{k}}$$

where $\mathrm{P}_\mathrm{s}^\mathrm{k}$ denotes $P_s$ associated with the triple t'= $\{x_i, x_j, y_k\}$. If $P^* \geq \delta_3 = 0.05/\binom{n+m}{3}$, the type of the merged trinet would be N or T. Otherwise, it is classified as S-type. The merged trinet T' contains the fixed pair $\{x_i, x_j\}$ and the new merged taxon $y^*$.

The next step is to determine both the type and the order of the merged trinet T'. If the merged trinet is classified as NT-type, we need to identify the most strongly supported split; Initialize $s(x_i x_j|y^*)=0$, where $s(x_i x_j|y^*)$ denotes the support score of the split $x_i x_j|y^*$; Initialize $s(x_i y^*|x_j)=0$, where $s(x_i\, y^*|x_j)$ denotes the support score of the split $x_i\, y^*|x_j$; Initialize $s(x_j\, y^*|x_i)=0$, where $s(x_j y^*|x_i)$ denotes the support score of the split $x_j y^*|x_i$. We iterate over all m trinets, if the trinet is NT $(x_i, x_j; y^*)$, $s(x_i x_j|y^*)= s(x_i x_j|y^*)-\log(P_f)$; if the trinet is NT $(x_i y^*|x_j)$, $s(x_i y^*|x_j)= s(x_i y^*|x_j)-\log(P_f)$; if the trinet is NT ($x_j$ $y^*|x_i$), $s(x_jy^*|x_i)$= $s(x_jy^*|x_i)$-log($P_f$). After processing all such trinets, we could determine the split that has the best support score. Without loss of generality, let $s(x_ix_j|y^*)$ be the highest value. The trinet T' is assigned to one of the types $N_2(x_i, x_j; y^*)$, $N_3(x_i; x_j, y^*)$ , $N_4(x_i; x_j; y^*)$ or $T_1$ ($x_i$, $x_j$, $y^*$) based on the average value of $|Lv_{1,1}$-$Lv_{1,2}|$ and $|Lv_{2,1}$-$Lv_{2,2}|$ from the associated trinets. We assign the multiplicative average of $P_s$, $P_f$ and w, the average of $|Lv_{1,1}$-$Lv_{1,2}|$ and $|Lv_{2,1}$-$Lv_{2,2}|$ from every trinet in t' to the merged trinet.

Conversely, if the merged trinet is classified as S-type, we identify the least supported split; suppose this split is $x_ix_j|y^*$. Initialize $ws(x_ix_j|y^*)=0$, where $ws(x_ix_j|y^*)$ denotes the support score of the split $x_ix_j$ | $y^*$; Initialize $ws(x_i\ y^*|x_j)=0$, where $ws(x_i\ y^*|xj)$ denotes the support score of the split $x_i$ $y^*|x_j$; Initialize $ws(x_j\ y^*|x_i)=0$, where $ws(x_j\ y^*|x_i)$ denotes the support score of the split $x_jy^*|x_i$. We iterate over all m trinets, if the trinet is $S_1(x_i, x_j; y^*)$ or $S_1(x_j, x_i; y^*)$ or $S_2(x_i; y^*; x_j)$ or $S_2(x_j; y^*; x_i)$ , $ws(x_ix_j|y^*)$= $ws(x_ix_j|y^*)$-log($P_s$), $ws(x_iy^*\ |\ x_j)$= $ws(x_iy^*\ |\ x_j)$+ log($P_s$), $ws(x_jy^*|x_i)$= $ws(x_jy^*|x_i)$+ log($P_s$); If the trinet is $S_1(x_i, y^*; x_j)$ or $S_1(y^*, x_i; x_j)$ or $S_2(x_i; x_j; y^*)$ or $S_2(y^*; x_j; x_i)$, $ws(x_ix_j|y^*)$= $ws(x_ix_j|y^*)$+log($P_s$), $ws(x_i\ y^*|x_j)$= $ws(x_iy^*|x_j)$- log($P_s$), $ws(x_jy^*|x_i)$= $ws(x_jy^*|\ x_i)$+ log($P_s$); If the trinet is $S_1(x_j, y^*; x_i)$ or $S_1(y^*, x_j; x_i)$ or $S_2(x_j; x_i; y^*)$ or $S_2(y^*; x_i; x_j)$ , $ws(x_ix_j|y^*)$= $ws(x_ix_j|y^*)$+log($P_s$), $ws(x_iy^*|x_j)$= $ws(x_iy^*|x_j)$+log($P_s$), $ws(x_jy^*|x_i)$= $ws(x_jy^*|x_i)$- log($P_s$). Without loss of generality, let $ws(x_ix_j|y^*)$ be the highest value.

We next need to assign a trinet whose worst split is $x_ix_j|y^*$. There are three possible trinets whose worst split is $x_ix_j|y^*$, that is $S_1(x_i, x_j; y^*)$, $S_2(x_i; y^*; x_j)$ and $S_2(x_j; y^*; x_i)$. Let $US_1=0$ be the updated support score of $S_1(x_i, x_j; y^*)$, $US_{21}=0$ be the updated support score of $S_2(x_i; y^*; x_j)$ and $US_{22}=0$ be the updated support score of $S_2(x_j; y^*; x_i)$. In this case, if the observed trinet is $S_1(x_i, x_j; y^*)$, $US_1$= $US_1$+(κ-$|Lv_{3,1}$-$Lv_{3,2}|$), $US_{21}$= $US_{21}$+($|Lv_{3,1}$-$Lv_{3,2}|$- κ), $US_{22}$= $US_{22}$+($|Lv_{3,1}$-$Lv_{3,2}|$- κ); If the observed trinet is $S_2(x_i; y^*; x_j)$, $US_1 = US_1 + (\kappa - |Lv_{3,1} - Lv_{3,2}|)$, $US_{21} = US_{21} + (|Lv_{3,1} - Lv_{3,2}| - \kappa)$, $US_{22} = US_{22} + (-|Lv_{3,1} - Lv_{3,2}| - \kappa)$; If the observed trinet is $S_2(x_j; y^*; x_i)$, $US_1 = US_1 + (\kappa - |Lv_{3,1} - Lv_{3,2}|)$, $US_{21} = US_{21} + (-|Lv_{3,1} - Lv_{3,2}| - \kappa)$, $US_{22} = US_{22} + (|Lv_{3,1} - Lv_{3,2}| - \kappa)$. After processing all such trinets, we could get the highest value in $US_1$, $US_{21}$ and $US_{22}$, and the corresponding trinet is what we choose to assign. We assign the multiplicative average of $P_s$, $P_f$ and w to the merged trinet, and the average of $|Lv_{3,1} - Lv_{3,2}|$ from every trinet in t' to the merged trinet.

### Threshold experiment

#### (i) Threshold experiment for distinguishing unimodal and bimodal signals in NT-type trinets.

As indicated by the heatmap in Fig. 2 in the main text, all N-type trinets exhibit highly consistent and monotonic behavior with respect to both the branch-length ratio and the number of loci. Therefore, it suffices to focus on a single representative trinet and assess whether an appropriate threshold can effectively distinguish unimodal and bimodal signals. In this study, we select $N_3$, where the cherry edge corresponds to a bimodal signal, whereas the pending edge represents a unimodal signal.

To determine an appropriate threshold κ for identifying bimodal signals in $N_3$, we conducted a threshold experiment under two ratios (ratio = 0.86 and 0.8) across varying numbers of loci. For each parameter setting, we performed 100 independent simulations. For cherry-type trinets, we recorded the 5th smallest value of $|Lv_{1,1} - Lv_{1,2}|$, whereas for pending-type trinets, we recorded the 5th largest value of $|Lv_{2,1} - Lv_{2,2}|$, providing conservative estimates for signal separation.

As shown in Table 1 and Table 2, both cherry and pending statistics increase monotonically with the number of loci, while cherry signals consistently grow at a substantially faster rate. We define the threshold as κ = number of loci / 125. Notably, even under the more challenging condition (ratio = 0.86), more than 60% of simulated data sets exhibit cherry statistics exceeding κ (Fig. 2), whereas all pending statistics remain below the threshold across all locus numbers considered.

These results are consistent with the heatmap patterns observed in the main text and indicate that κ = number of loci / 125 (corresponding to κ = 8 when loci = 1000) provides a reliable and robust criterion for distinguishing bimodal cherry signals from unimodal pending signals in $N_3$ trinets.

| **N3 (ratio = 0.86)** | **500** | **1000** | **2000** | **3000** | **4000** | **5000** | **6000** | **7000** | **8000** | **9000** | **10000** |
|---|---|---|---|---|---|---|---|---|---|---|---|
| **Cherry** | 0.77 | 2.69 | 9.36 | 13.86 | 20.16 | 25.02 | 30.18 | 37.79 | 44.07 | 54.59 | 59.65 |
| **Pending** | 4.89 | 5.31 | 8.33 | 14.63 | 15.16 | 15.42 | 19.17 | 20.55 | 26.39 | 25.51 | 27.91 |
| **Average** | 2.83 | 4.00 | 8.80 | 14.20 | 17.60 | 20.10 | 24.70 | 29.20 | 35.23 | 40.00 | 43.50 |

Table 1 Threshold experiment ($N_3$, ratio = 0.86)

| **N3 (ratio = 0.8)** | **500** | **1000** | **2000** | **3000** | **4000** | **5000** | **6000** | **7000** | **8000** | **9000** | **10000** |
|---|---|---|---|---|---|---|---|---|---|---|---|
| **Cherry** | 8.93 | 24.58 | 52.94 | 82.37 | 114.47 | 145.06 | 190.19 | 219.17 | 263.20 | 293.41 | 319.62 |
| **Pending** | 3.73 | 5.55 | 10.50 | 15.10 | 15.63 | 17.67 | 22.98 | 23.44 | 23.40 | 29.03 | 35.07 |
| **Average** | 6.33 | 15.07 | 31.72 | 49.00 | 66.07 | 81.36 | 107.45 | 121.30 | 149.22 | 161.22 | 177.34 |

Table 2 Threshold experiment ($N_3$, ratio = 0.8)

#### (ii) Threshold experiment for distinguishing $S_1$ and $S_2$ trinets.

To further assess the applicability of the threshold κ = number of loci / 125 to S-type trinets, we examined its performance under a challenging setting with ratio = 0.9 and number of loci = 1600. Figure 2 shows the distribution of $|Lv_{3,1}-Lv_{3,2}|$ across 100 simulation replicates, where the left panel corresponds to S1 and the right panel corresponds to $S_2$.

Under this setting, only 3 out of 100 $S_1$ replicates exceed the threshold κ, whereas all $S_2$ replicates lie above the threshold. This clear separation indicates that κ = number of loci / 125 remains effective for distinguishing $S_1$ and $S_2$ even when the branch-length ratio is 0.9.

Compared with NT-type trinets, S-type trinets are generally easier to detect in terms of bimodality. This is because, for S-type trinets, two sets of parameters are optimized independently using L-BFGS-B according to the split distributions, whereas for NT-type trinets, only a single parameter set can be optimized. As a result, bimodal signals in S-type trinets are amplified more effectively, leading to stronger separation between unimodal ($S_1$) and bimodal ($S_2$) cases.

Taken together, these results demonstrate that even at ratio = 0.9, the threshold κ = number of loci / 125 provides a robust and reliable criterion for distinguishing $S_1$ and $S_2$ trinets.

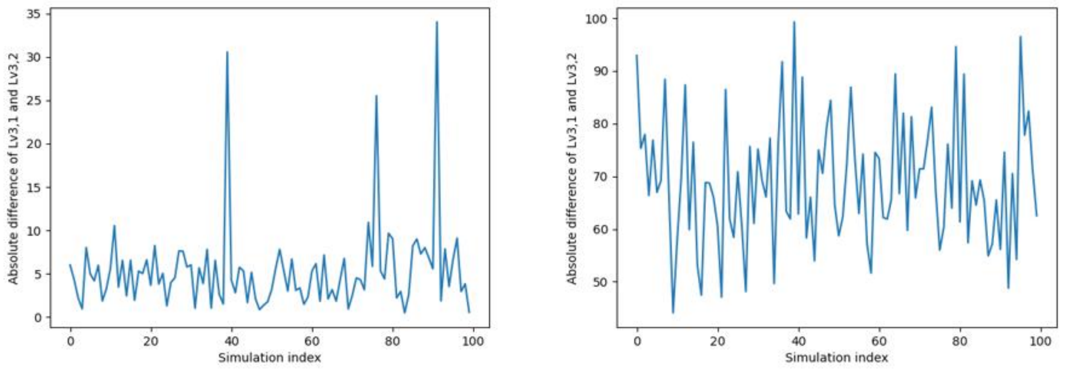


i ii

**FIG. 2. Distribution of $|Lv_{3,1}-Lv_{3,2}|$ across 100 replicates for $S_1$(i) and $S_2$(ii) trinets (number of loci = 1600, ratio=0.9)**
